# Quantitative analysis of proteomic changes in monoclonal MDCK cell lines infected with human influenza A virus

**DOI:** 10.1101/2025.06.25.661656

**Authors:** Jan Küchler, Tilia Zinnecker, Patrick Hellwig, Maximilian Wolf, Dirk Benndorf, Yvonne Genzel, Udo Reichl

## Abstract

Suspension MDCK cells are a substrate for producing influenza A virus (IAV) and typically show very high virus yields compared to other animal cells. Due to the significant heterogeneity within cell populations, studying and comparing clonal cell lines with regard to specific properties, such as superior growth or higher productivity, could facilitate process optimization. In this study, we analyzed the expressed proteins of clonal cell lines to identify intrinsic characteristics of effective IAV producers. We compared proteome changes in two IAV-infected monoclonal suspension MDCK cell lines: C59, a low-yield IAV producer with fast cell growth and small cell diameter, and C113, a high-yield IAV producer with average cell growth and large cell diameter. We examined growth rate, size, metabolism and IAV production.

A total of 5177 host cell proteins were detected in both cell lines using DIA-PASEF mode with a TimsTOFpro mass spectrometer. Analysis of the differentially expressed proteins revealed that fatty acid oxidation and branched-chain amino acid degradation were upregulated in highly productive cells. In contrast, steroid biosynthesis and DNA replication were more active in faster-growing cells. Following infection, 122 proteins were significantly upregulated (p<0.05, log_2_-fold change ≥1) in the high-producing cell line. These proteins were associated with membrane trafficking, interactions with the IAV-NS1 protein and virus production. Additionally, 98 proteins associated with antiviral pathways such as the proto-oncogenic receptor tyrosine kinase MET and tumor necrosis factor (TNF) signaling were downregulated (p<0.05, log_2_-fold change ≤1).

In the cell line that produced lower IAV titers, 77 proteins were downregulated and 57 were upregulated after infection. RNA metabolism appeared to be downregulated, while the tricarboxylic acid (TCA) cycle and the stress response were both upregulated. In the high-yield C113 clone, only proteins associated with apoptosis and the target of rapamycin kinase (TOR) were expressed following infection. This may indicate a more effective release of virus particles. A comparison of intracellular IAV protein levels demonstrated that M1 and NA levels were 4-fold and 8-fold higher, respectively, for the high-yield C113 cell line. These findings again suggest better virus release.

## Introduction

Most influenza vaccines are still produced using embryonated chicken eggs. However, animal cell culture technology offers several advantages over egg-based production. These advantages include faster production times during pandemics, improved scalability, enhanced immunogenicity, better process control, and the ability to operate in closed systems [1]-[5].

Vero cells and Madin-Darby canine kidney cells (MDCK) are among the most promising candidates for the cell culture-based production of influenza viruses [6]-[7]. MDCK cells are used to produce Flucelvax Tetra (CSL Seqirus), for example, and are generally considered as a preferred substrate due to their high susceptibility and yields for a wide range of influenza viruses [3]-[4] [8]-[12]. Additionally, suspension cell lines offer several advantages for industrial viral vaccine production because they can be grown to high densities, enabling large-scale production. Furthermore, chemically-defined media that are free of animal components are available [5] [12]-[13].

From work on CHO cells, it is known that there are significant differences in cell lines due to the passaging and cultivation history. When switching to cell line development and generation of cell clones, clear differences are observed between clones and cell populations. Selecting specific cell clones for specific needs has become standard for recombinant protein production. However, this is not yet widely considered for virus production [4]-[5]. MDCK cell populations are also described as highly variable with respect to their growth properties, metabolism, virus titer, antigen glycosylation patterns and morphology [13]-[17]. Due to this inherent heterogeneity, clone selection can be leveraged for effective influenza A virus (IAV) production [6]. To analyze and compare the properties of clonal cell lines, proteomic analysis via mass spectrometry (MS) can be applied. Compared to other analytical methods, proteomics uniquely captures the cellular status, activated signaling pathways and metabolic shifts inside the cell during IAV infection. Quantitative MS, in particular, allows for an in-depth analysis of proteins produced during cell growth and IAV replication. This enables a comprehensive comparison of different cell lines [18]-[21]. Thus far, we have employed this approach to compare the IAV proteins produced by different cell lines cultivated in different media [22].

In this study, we use our previously described proteomic workflow for IAV protein production and expand it to cover the proteome of two IAV-infected clonal MDCK cell lines. As previously reported, clones C59 and C113 exhibit distinct characteristics regarding growth, metabolism, and virus production [6]. To improve our understanding of the high-yield clone C113, we analyzed these two different cell lines at the proteomic level. Our goal was to track IAV protein production and identify key host cell factors and metabolic pathways associated with high IAV titers in MDCK cells. Using this approach, we detected over 5000 host cell proteins in both cell lines and mapped them to several hundred metabolic pathways. This allowed us to compare the two cell lines comprehensively. Despite their origin from the same parental cell line and growth under identical cell culture conditions, our results demonstrate significant differences between the two monoclonal MDCK cell lines.

## Materials and methods

### Cells and viruses

The two MDCK clones C59 and C113, developed and provided by Sartorius (Germany) and characterized in previous studies [6], were used for all experiments. These two clones were created through single-cell cloning from adherent MDCK cells (#CCL-34, ATCC) and were adapted to suspension growth [6]. The monoclonal cell lines were cultivated in 4Cell® MDXK CD medium (Sartorius, Germany), supplemented with 8 mM L-glutamine. Cells were maintained in non-baffled shake flasks (Corning, #431143) with a working volume of 30 mL and cultivated at 37°C and 5% CO2, with a shaking frequency of 120 rpm (50 mm throw) in a Multitron Incubator shaker (50 mm shaking orbit, Infors AG, Switzerland). Passaging was performed 2-3 times per week, and cells were inoculated at viable cell concentrations (VCCs) ranging from 2-8 × 1E+05 cells/mL. Cell viability, diameter, and concentration were measured using a Vicell XR automated cell counter (Beckman Coulter, #731050).

For infection, influenza A/PR/8/34 (H1N1, RKI, 3.0 log_10_ (HA units/100 µl), 1.10E+09 TCID_50_/mL) seed virus adapted to adherent MDCK cells (#84121903, ECACC) was used. Virus stock was diluted to achieve a multiplicity of infection (MOI) of 3. Trypsin (Thermo Fisher Scientific, no. 27250-018) was added at time of infection (final activity of 20 U/mL). A medium exchange was performed 1 h post infection (hpi) to remove remaining seed virus. Samples were taken before infection (mock) or at 12 hpi based on preliminary experiments.

### Analytics

Virus production was monitored using the hemagglutination (HA) [21] and TCID_50_ assays to determine the total and infectious number of virions produced, respectively. For both assays, samples were centrifuged (3000 x g) and the virus-containing supernatant was collected and stored at -80°C before analysis. Total virus particle concentrations (C_vir_) were estimated using the erythrocyte concentration (C_ery_) in the HA assay. The standard deviation of the HA assay was ±0.081 log10 (HA units/100 µL). Infectious virus particle concentrations were determined using a 50% tissue culture infectious dose (TCID_50_) assay, as described by Genzel and Reichl [21]. In this assay, adherent MDCK cells were infected with serial dilutions of the virus-containing samples in the presence of 50 units/mL trypsin. The cells were stained with an HA-specific antibody 48 hpi. The dilution error of the TCID_50_ assay was 0.3 log, as reported by Genzel and Reichl [21].

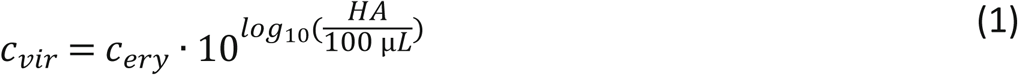

### Proteomics workflow and peptide quantification

Five IAV proteins (hemagglutinin (HA), nucleoprotein (NP), neuraminidase (NA), matrix protein 1 (M1) and non-structural protein 1 (NS1)) were quantified over a single infection cycle using absolute protein quantification (AQUA) as described by Küchler et al. 2025 [22] (Fig 1). A minimum of three AQUA peptides were used for quantification of each protein. Infected cells were sampled at 12 hpi and processed as follows: cell pellets were washed once with 1 mL PBS, then lysed using RIPA buffer (Thermo Fisher Scientific, Waltham, Massachusetts, USA) and 26G needles (Terumo Agani, Tokio, Japan). For approximately 4 × 1E+06 cells, 300 µL of lysis buffer was used. Protein concentrations were determined using a commercial BCA assay (Thermo Fisher Scientific, Waltham, Massachusetts, USA), and 50 µg of each sample was used for further processing and preparation for MS measurements using a modified filter-aided sample preparation (FASP) method [23].

**Figure 1:**
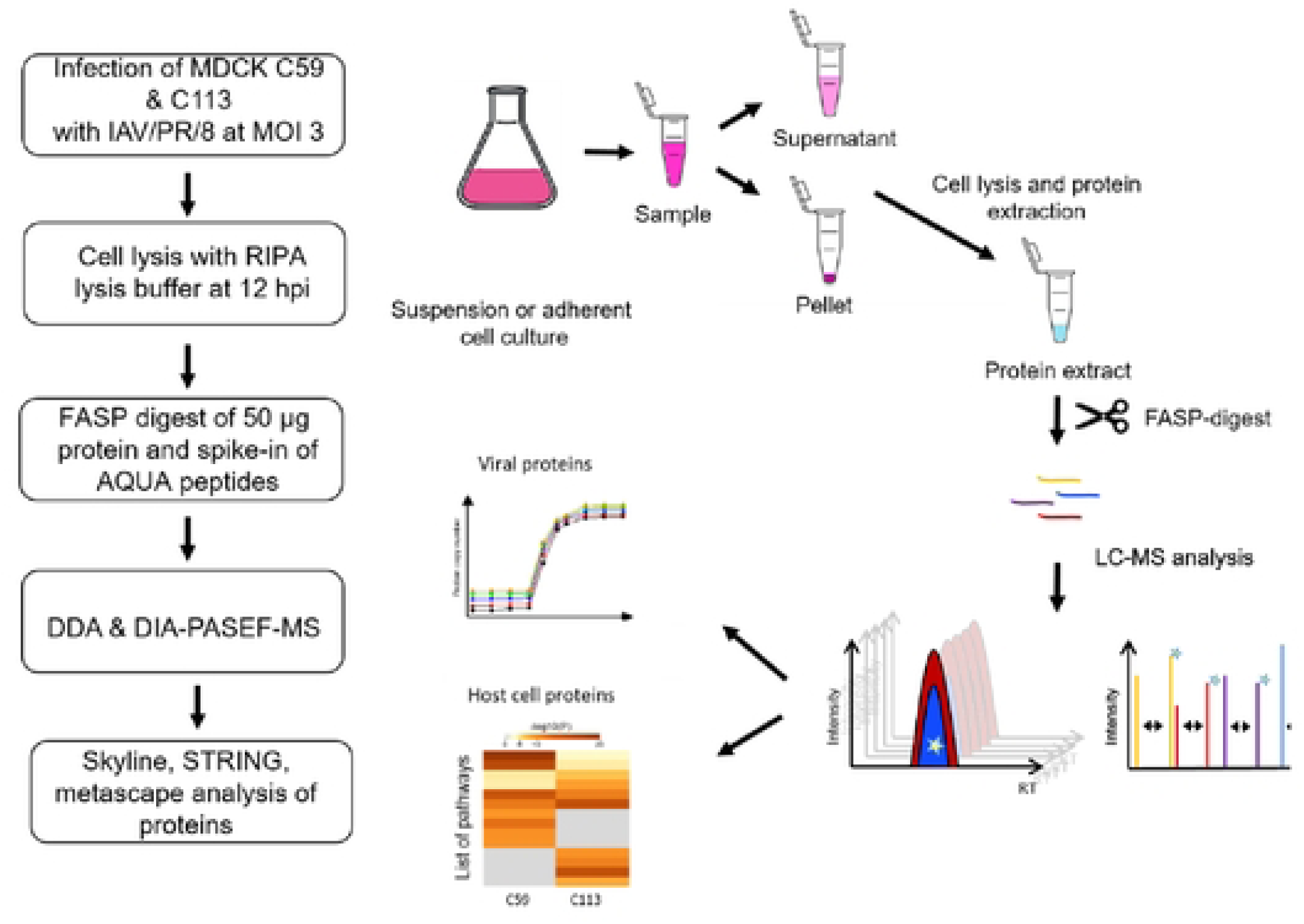
Workflow for the proteomic study comparing CS9 and Cl13MOCK cellclones and their IAV production. RIPA: radio­ **immunoprecipitation assay; FASP: filter·aided sample preparation; AQUA: absolute quantification; ODA: data-dependent** acquisition; DIA:data-independent acquisition; PASEF: parallel accumulation serial fragmentation. (Adapted from [221).

**Figure 2:**
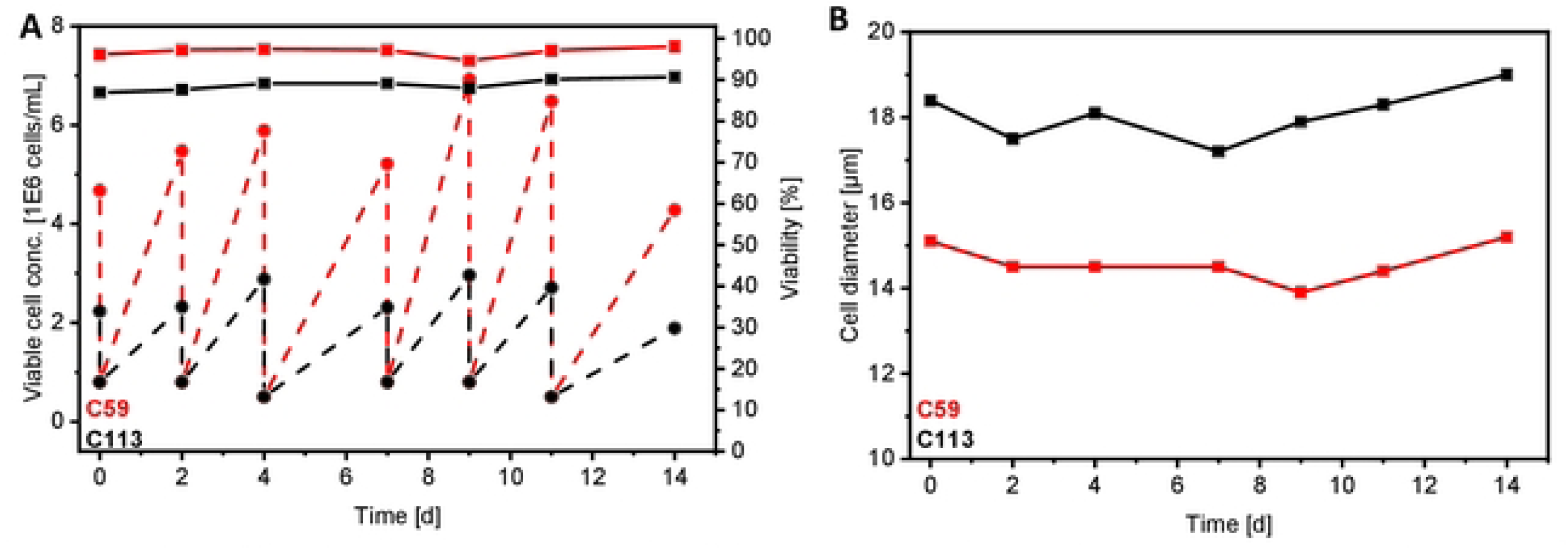
Cell growth of the two monoclonal MOCK cell lines (C59: red, Cll3: black) over several passages. A: viable cell concentration (dashed line) and viability (solid line). B: cell diameter. Both cell lines were seeded at a constant cell **concentration of 5·8 1E6 cells/ml, with a seeding interval of 2·3 days.**

Proteolytic digestion was performed using 2.5 µg (1:20) of MS-approved trypsin for 16 hours at 37°C. Dried samples were then diluted in 90 µL of load A buffer (LC-MS grade water + 0.1% trifluoric acid) plus 10 µL of AQUA standard mix (0.2 pmol/µL of each peptide). AQUA peptides with isotopic labels were used to quantify IAV peptides by back-calculating to the absolute copy number after adding a specific amount to each sample. The AQUA standard mixture contained 20 peptides for the five major IAV proteins with a heavy label (all C and N atoms of the last amino acid, either lysine or arginine, were replaced with their C13 or N15 isotopes) (Thermo Fisher Scientific, Waltham, Massachusetts, USA). For each sample, 2 µL (1 µg protein, 0.04 pmol AQUA peptides) were injected into the LC-MS/MS system (see below), enabling the quantification of approximately 1.0E+08 protein copies/cell using the Skyline software, as determined by previous experiments and mathematical modelling approaches [22] [24].

Liquid chromatography (LC) and MS measurements were performed using an UltiMate 3000 system (Thermo Fisher Scientific, Waltham, Massachusetts, USA) and a TimsTOFpro (Bruker Daltonik, Bremen, Germany). The MS was operated in data-dependent (DDA) Parallel Accumulation Serial Fragmentation (PASEF) mode and in data-independent (DIA)-PASEF mode as described before [25]. Windows for DIA-PASEF scans were determined based on DDA measurements using py_diAID (version 0.030) as described in Skowronek et al. [25] resulting in a 99.80 % precursor coverage (Supplementary figure S1). A spectrum library was created and the raw data analyzed with the fragpipe software (version 22.0) and DIA-NN software (version 2.0.2) as described before [26]. The reference genome used for the spectrum library was “*C. l. familiaris*” downloaded from Uniprot (5/9/2025). The resulting DIA-NN files (.parquet) were filtered by q-value (Q.value and Global.Q.Value) and a pivot table was created in R Studio (version 2023.06.2 build 561) using an in-house script. The protein groups were functionally annotated using Prophane [27].Subsequent comparison of relative protein expression between the cell lines was performed in Excel; significance was tested by using a t-test (p<0.05, two-tailed, unequal variances). All detected proteins and their abundancies, as well as those visualized in volcano plots and heatmaps are listed in Supplementary file 2. Principal coordinate analysis (PCoA) was performed and visualized with Python (Supplementary file 1) using Numpy, (1.26.4), Scikit-learn (1.4.2), Scipy (1.13.0) and matplotlib (3.8.4). For protein network analysis, we used the online tool STRING (string-db.org; [28]) with *H. sapiens* or *C. l. familiaris*. Venn diagrams, volcano plots and heat maps were created using the respective add-ins in OriginPro (version 9.6.0.172). Protein expressed only in the infected or mock samples were analyzed using the Metascape tool (metascape.org, [29]).

## Results and discussion

### Growth characteristics and virus production of two different monoclonal cell lines

First, the cell growth and virus production of the two monoclonal MDCK cell lines, C59 and C113, were compared to confirm previously observed differences [6]. The C59 clone exhibited faster cell growth, achieving a viable cell concentration that was 2-3 times higher at each passage, with high viability (Fig 1A). The diameter of the C59 cell clone was on average about 3-4 µm smaller compared to C113 (Fig 1B), which was in line with previous observations [6]. Assuming the cells were spheres, this resulted in a 1.6-fold surface area difference and a 2-fold volume difference. However, both are within the normal range for MDCK cells [15]. Note: for these calculations, we assumed a smooth surface. If there are a lot of microvilli, the real surface area could be much larger and the differences between the cell lines more pronounced. This would affect the metabolism, IAV budding and virion release [30]-[32].

Furthermore, the total (HA assay) and infectious (TCID_50_) virus titers were determined for both cell lines (Table 1). The clone C113 produced 0.9 log_10_(HAU/100 µL) more virions and about 9-fold more infectious IAV, which is consistent with earlier reports [6]. Together with the higher VCCmax of the C59 clone, this results in a significantly higher cell-specific virus yield for C113. When the difference in cell size is taken into account (volume-specific virus yield), a substantially higher viral yield is also observed for C113 (Table 1). This demonstrates that, in addition to cell size, intrinsic factors may also contribute to the difference in virus yield as it was also shown in a previous study on MDCK cells [15]. Moreover, the C113 clone produces higher proportions of infectious virions than C59 (smaller ratio of CSVY_HA_/CSVY_TCID_). Overall, compared to our previous studies, the values obtained in this study are within a similar range [4] 6].

**Table 1:**
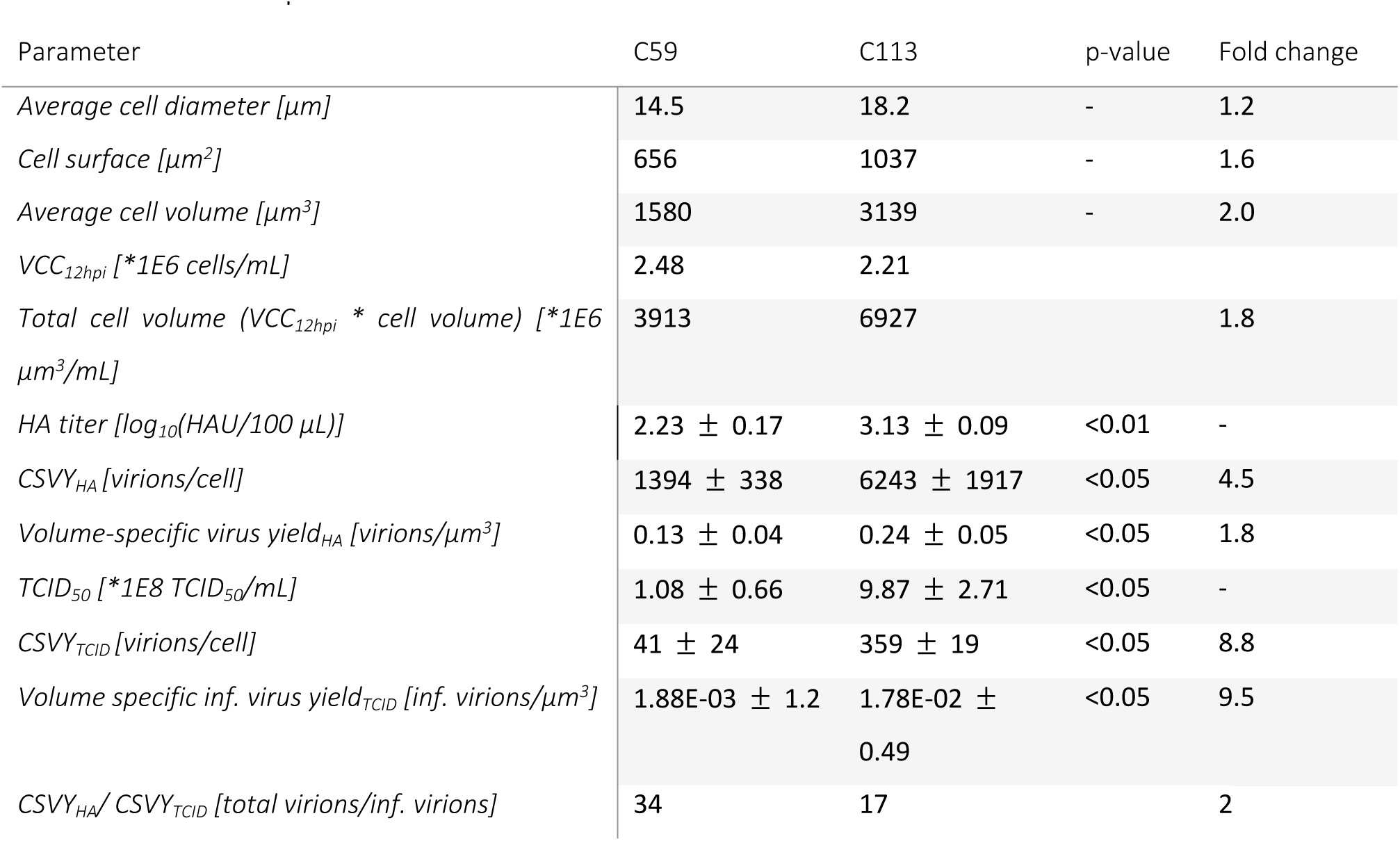
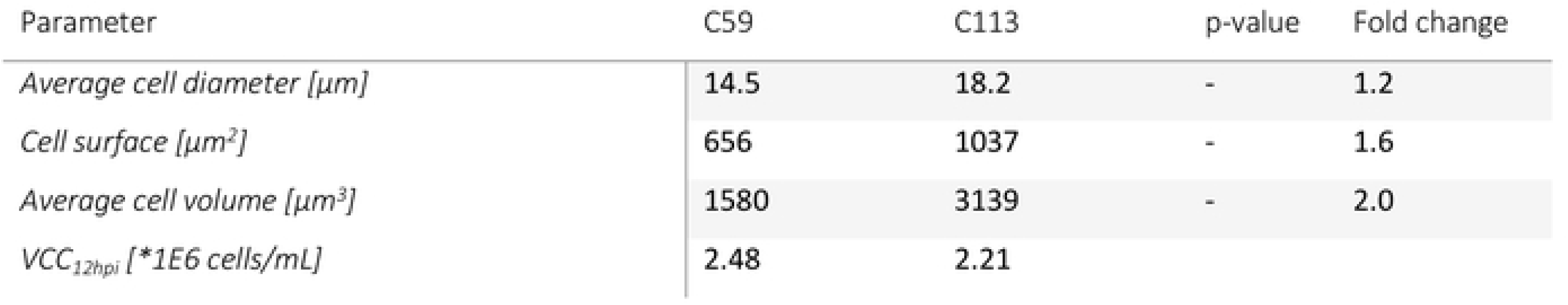
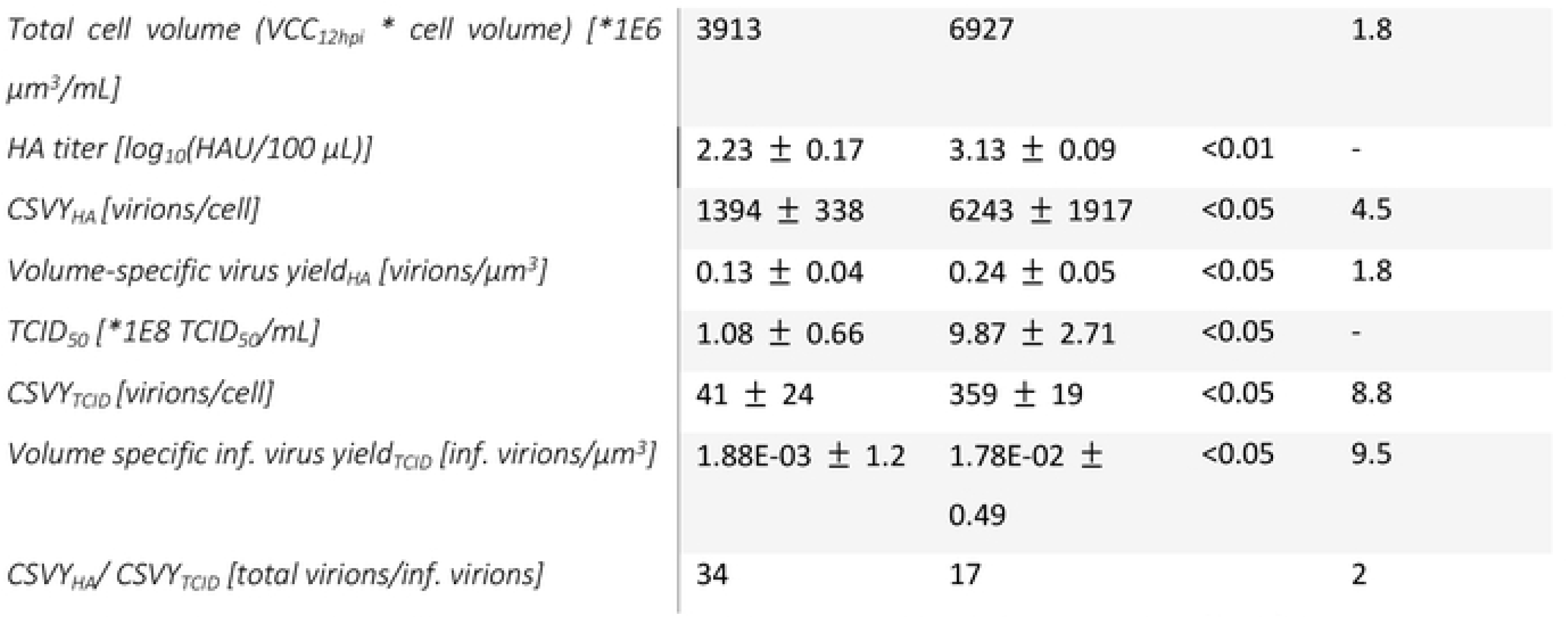
IAV production of the monoclonal MOCK celllines CS9 andC113.

### Influenza A virus protein production of the two different monoclonal cell lines

In order to compare the two monoclonal cell lines on a molecular level, the intracellular IAV protein production was analyzed. For this, the absolute protein copy numbers of the five most abundant IAV proteins during the infection process were determined by employing a previously published assay from our group [22]. As shown in Figure 3, the majority of IAV proteins were produced at comparable intracellular levels by both cell lines, which is consistent with previously published findings [22]. The concentrations of HA, NP and NS1 proteins were almost identical for both cell lines with less than 2- fold difference, mostly with slightly higher concentrations for the C113 clone. Nevertheless, a 4-fold increase in M1 protein concentrations and a 8-fold increase in NA protein concentration was observed in the C113 clone. The higher concentrations of NA protein observed in C113 could potentially favor virus assembly and budding since both proteins play a critical in the efficient release of progeny virions [33]. Furthermore, the ratio of HA to NA protein (HA/NA balance) plays a very important role in the virus production [34]. The higher copy numbers of HA compared to NA in the C59 clone could lower the rate of virus release. Furthermore, the M1 protein plays a pivotal role in virus budding. It is associated with the HA and NA proteins and it binds to the viral RNA complexes [35].

**Figure 3:**
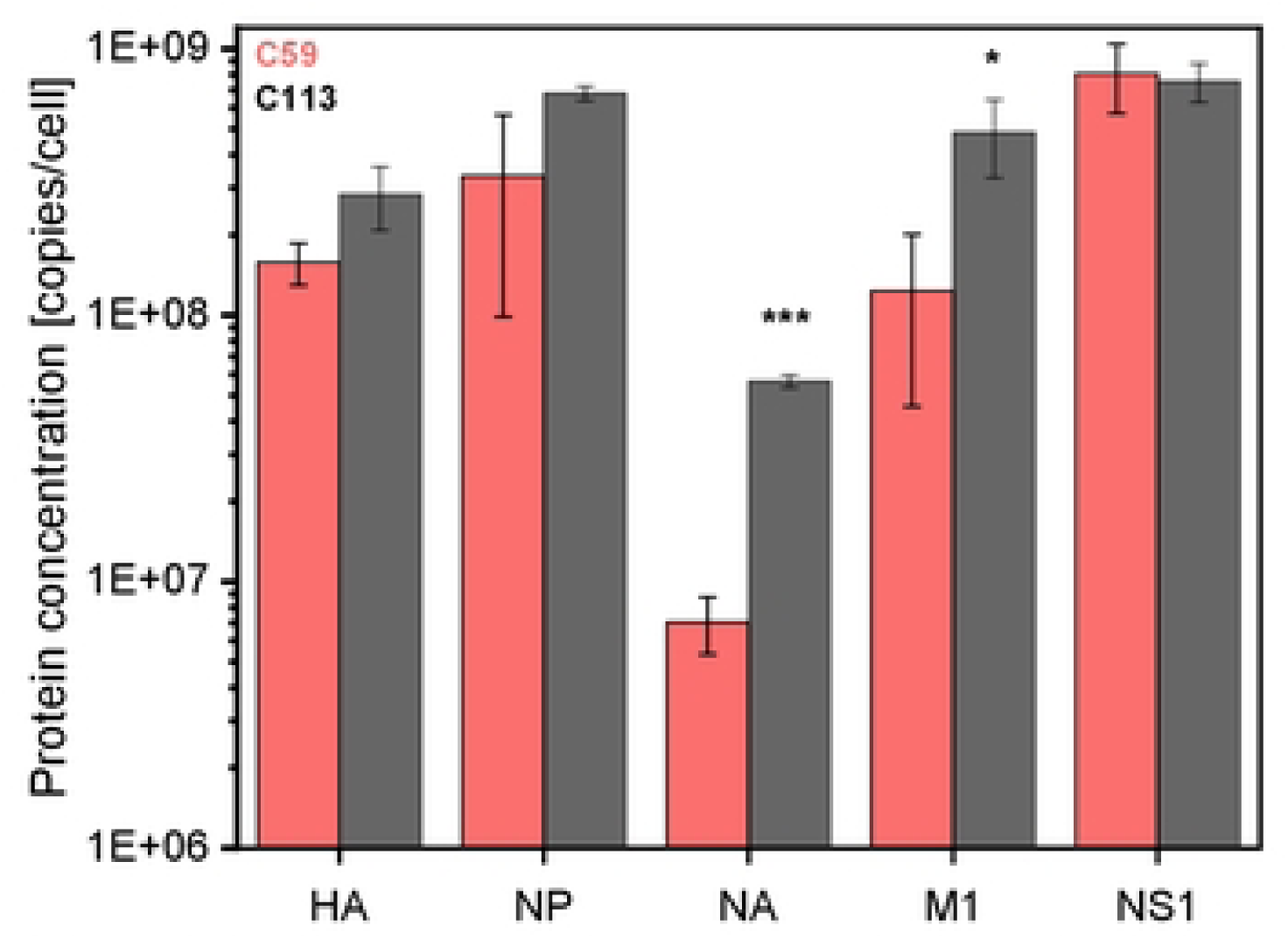
Intracellular IAV protein production of the monoclonal MOCK cell lines C59 (red) andC113 (grey). Absolute protein copynumbers for fiveIAV proteins at 12 hpi were determined based on isotope-labelled peptides (22).Cells were seeded in trypsin-containingMOXK mediumin shake flasks with a workingvolume of 30 ml, and directly infected at 2.0E+06 cells/ml with A/PR/8/34 (HlNl, RKI) with an MOI of 3. Samples were taken at 12 hpi. A complete medium exchange was performed at l hpi. Selected peptides were quantified by adding a defined amount of AQUA standard peptides into each sample using DIA-PASEF-MS, and analyzed with theSkyline software. A list of the labelled peptides including the limit of detection and the **limit of quantification is provided in the Supplementary tables S1, S2 and S3. Significance was tested with a two-sided** student’s !•test (*p<0.05, •••p<0.001).

### Host cell protein analysis of the two different monoclonal cell lines

First, differences in host cell protein expression, irrespective of their relative abundances were compared between the cell lines. Therefore, DIA-PASEF measurements of the samples were analyzed with *C. l. familiaris* proteome using fragpipe and DIA-NN to create a spectral library. The latter was then processed with an R script (see Supplementary file 3) and the online tool Prophane.

As an initial step a principal coordinate analysis (PCoA) was performed (Supplementary file 1). As illustrated in Figure 4A, the three replicates of all four sample sets are closely grouped, indicating that the two cell lines and the two cell states (mock versus infected) exhibit distinct patterns of protein expression. Interestingly, for C113, protein expression of the three infected replicates seems to be more similar to each other compared to the infected replicates of C59 (Fig 4A).

**Figure 4:**
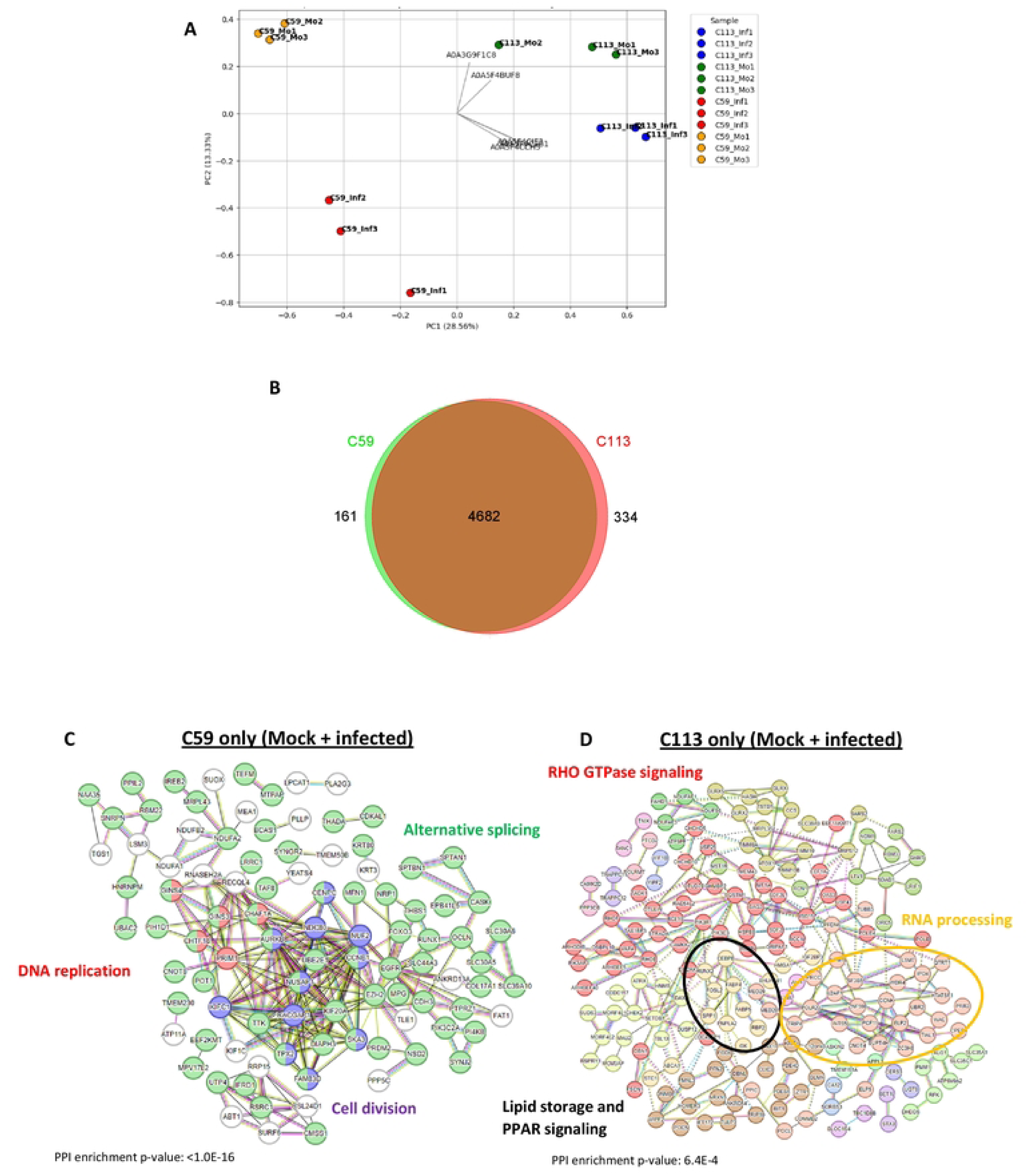
Comparison of protein expression of the two monoclonal MOCK cell lines CS9 and Cll3. A:principal coordinate analysis of all detected proteins across the three replicates of CS9 infected (CS9_1nf), CS9 mock (CS9_Mo), C113 infected (C113_1nf) and Cll3 mock (Cll3_Mo) using Bray-Curtis distance (Supplemental file 1). 8: Venn diagram of all detected proteins (C59 andC113) in at least two of the three replicatesper condition. C,D: proteinnetwork anatysisof proteins$which are only expressed by one cell clone using the STRING platform and *H. sapiens* as the reference proteome. Each protein is assigned to the corresponding gene. Colors refer to functional clusters, which are connected with straight line, genes of different clusters with a functional similarity are connected with a dashed line.Genes that belong to two functional clusters

Across all samples, 5507 proteins were identified, 5177 of which could be detected in at least two replicates of the C59 or C113 clones. Of the 5177 proteins, about 6.5 % (334) were exclusively detected in C113, 90 % (4682) in both cell lines and 3.5 % (161) in C59 only (Fig 4B) (Supplementary file 2).

The proteins, which were detected in only one of the two cell clones, were subsequently analyzed using the protein network platform STRING. *H. sapiens* was selected as the host due to the more comprehensive information available about the proteins. However, we ensured that each gene name was annotated to the same protein for both *C. l. familiaris* and *H. sapiens*. As shown in Supplementary figure 2, for *H. sapiens,* more information regarding cross-connections between proteins is available, resulting in a more interconnected network. Moreover, for all comparisons of Figure 4, protein-protein interaction enriched p-values were below 0.05, indicating that the connections between the proteins were not random (string-db.org). For proteins detected exclusively in the C59 clone, the majority were associated with alternative splicing (Fig 4C). In contrast, proteins detected only in the C113 clone, could be functionally assigned to Ras homologue (RHO) GTPase signaling and lipid storage (Fig 4D). Alternative splicing was been demonstrated to play a pivotal role in the proliferation of cancer cells. This process enhances the efficiency of the energy metabolism and can prevent apoptosis [36]. In addition, RHO GTPase signaling is a key regulator of actin and microtubule dynamics and can influence numerous processes in cancer cells such as membrane trafficking, cell cycle and Phosphoinosit 3- Kinase (PI3K) signaling. Furthermore, it can support viral processes during infection [37]-[38].

In the next step, we compared differences in the amount of the expressed proteins the sample sets (Fig 5A-D), i.e., samples without infection (mock) and after infection. For the mock samples, 266 of the 5177 detected proteins in both cell lines were significantly different expressed (119 upregulated in C59, 147 upregulated in C113). With a 40-fold increase in expression in the C113 clone, the succinyl-CoA:3-ketoacid-coenzyme A transferase (Acc: A0A8I3MS17, Gene: OXCT1) was the most upregulated protein compared to C59. This protein is a key enzyme for ketone metabolism, sterol production, tumorigenesis and signaling in tissues and cancer cells [40]-[41]. In relation to that finding, higher levels of sterols were found in the C113 clone compared to C59. In the case of the C59 mock infection, four of the proteins that were upregulated were in a similar range with an about 16-18 fold higher abundancy than in the C113 clone. Two of these are classified as protein-disulfide isomerases which have been shown to have a positive effect on the proliferation of cancer cells (Acc: A0A8I3Q0F7, Gene: PDIA6, Bai et al., 2019; Acc: A0A8I3MZE2, Gene; CRELD1, [42]), one as an aminopeptidase with antiviral effects (Acc: A0A8I3PTW1, Gene: LTA4H, [43]) and one as a B-cell receptor-associated protein, for which the function has not yet been described (Acc: A0A8I3PYE6, Gene: BAP29, [44]).

**Figure 5:**
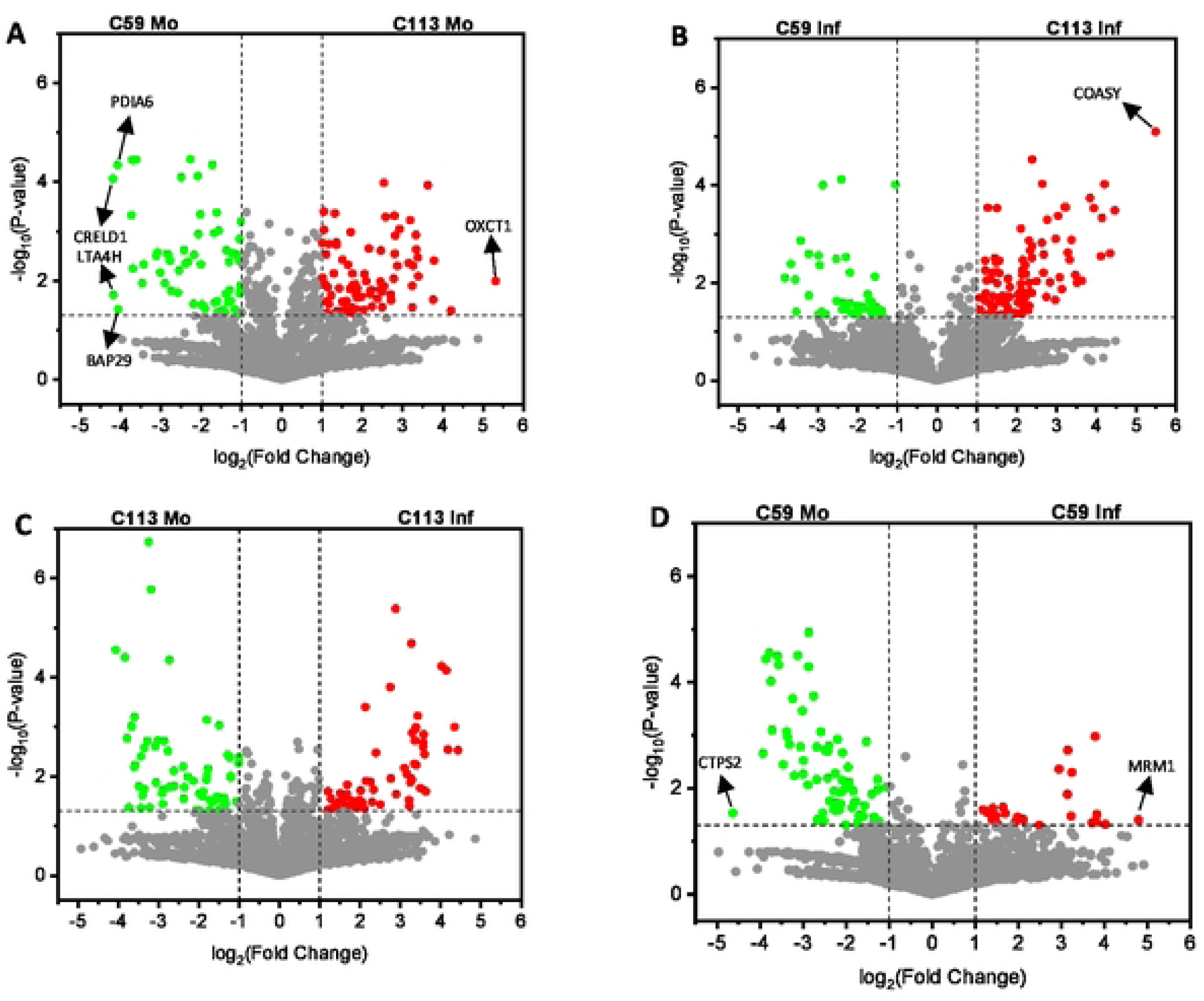
C**o**mparison **of protein expression profiles of the two monoclonal MOCK celllines CS9 and C113 without infection** (Mo) and after infection (Inf). A-0: volcano plots; for the calculation of the fold-change, average ratios of mock (Mo) or infected (Inf) were calculated for all proteins detected in all replicates, respectively. Each point corresponds to oneprotein. Green andredproteins were significantly different expressed (p-value <0.05) witha fold change of >2 (grey lines). Statistical significance wascalculated with student’s t test (two-sided, heterogeneous variances).

For the infected samples, 179 proteins were found to be significantly upregulated in C59 (57) or C113 (122) (Fig 5B). Again, for the C113 clone, the bifunctional coenzyme A synthase, exhibited a 45-fold increase in expression compared to C59. The bifunctional coenzyme A synthase (Acc: A0A8I3N4F7, Gene: COASY) is involved in the cofactor biosynthesis for multiple lipids [45]. Interestingly, this protein has been described as a positive regulator of PI3K signaling [46], a process that has been shown to promote IAV infection during both early and late stages [47].

A comparative analysis of the changes after infection of each cell line individually (Fig 5C, D) revealed that for the C113 clone three times more proteins were significantly upregulated compared to C59 (100 vs. 31). In contrast, the number of downregulated proteins showed more similarity (94 vs. 76). For the C113 clone, multiple proteins were either up- or downregulated during infection. For C59, two proteins were identified with very high changes in expression levels: the cytidine triphosphate (CTP) synthase 2 (Acc: A0A8I3QAX6, Gene: CTPS2) was downregulated 25-fold, whereas the rRNA methyltransferase 1 (Acc: A0A8I3Q638, Gene: MRM1) was upregulated 28-fold. CTPS2 is associated with cell proliferation [48], whereas MRM1 plays a crucial role in RNA modification [49].

Subsequently, a protein network analysis was performed on the significantly differentially expressed proteins (student’s t-test, p<0.05; log2 fold changes >1 and <-1) to identify patterns and cross connections. To this end, all significantly differently expressed proteins from each comparison (C113Mo vs. C59Mo, C113Inf vs. C59Inf) were analyzed using the functional protein expression network tool STRING (reference proteome: *H. sapiens*). The protein networks and related pathways are shown in the Supplementary (Figure 2).

A comparison of the protein networks of the two cell clones without infection (mock), reveals differences in fatty acid metabolism and DNA replication (Fig 6A, B, Supplementary figure 2). While the steroid biosynthesis was found to be more active in the C59 clone compared to C113, the fatty acid beta oxidation and the degradation of specific amino acids was upregulated in C113. Among the various steroids, cholesterol has been identified as a significant player. Cholesterol biosynthesis has been demonstrated to be beneficial for the proliferation of cancer cells and the replication of IAV, where it plays a pivotal role for cell survival and viral budding [50]-[53]. Other steroids are also described to act as antivirals inside the cell [54]. The second group of proteins, which was found to be upregulated in the uninfected C59 cells (mock) is related to DNA replication and proliferation. The majority of these proteins have been found to be associated with cell cycle progression and mitosis. Therefore, the faster growth of the C59 clone may be ascribed to this observation given that these proteins promote cell proliferation [55].

**Figure 6:**
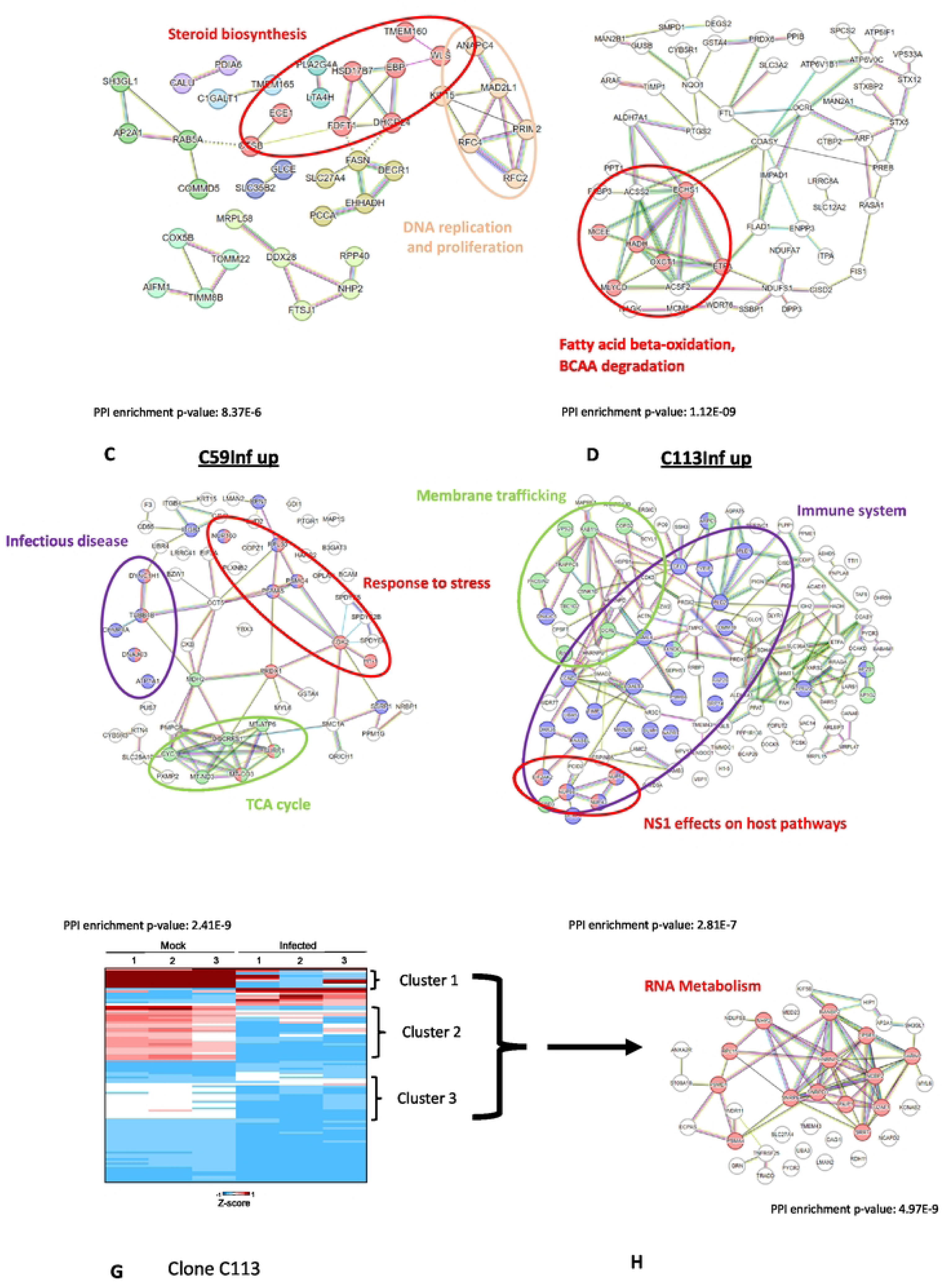
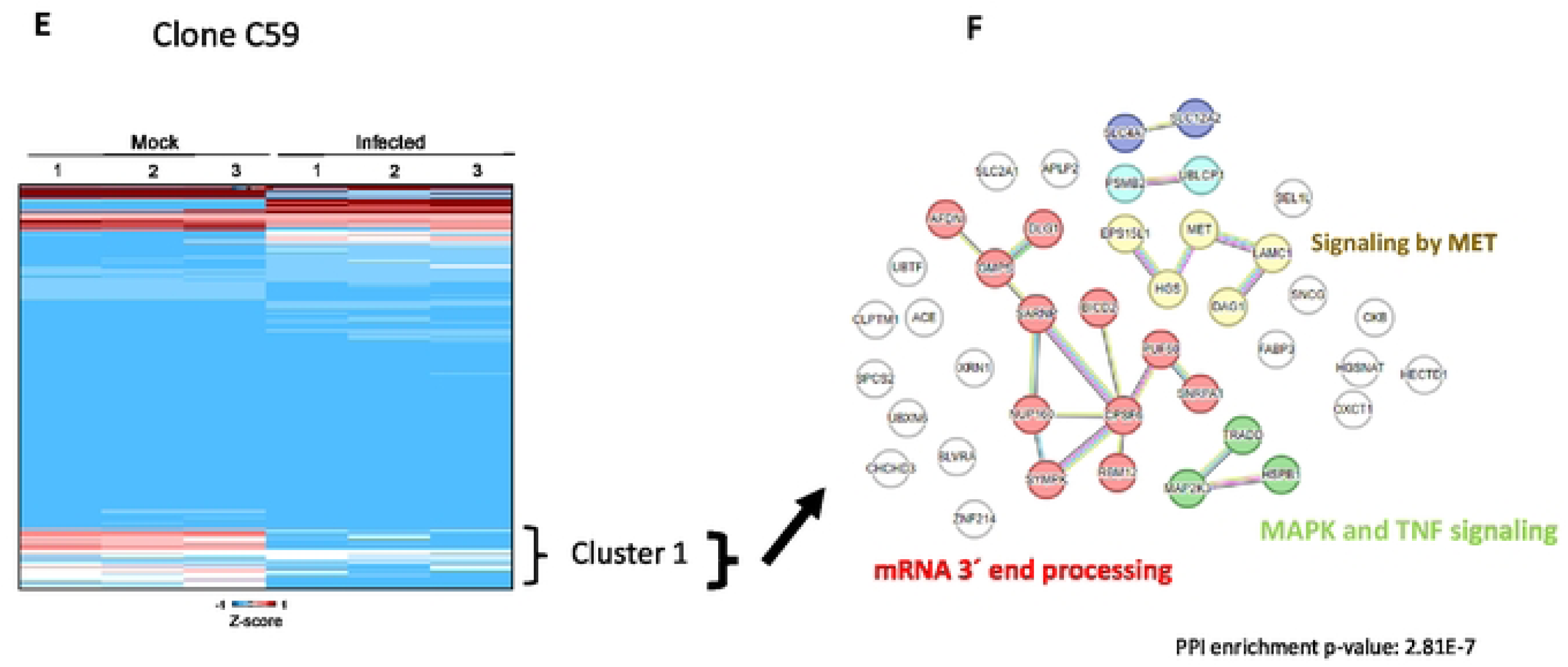
Analysis of significantly differentiallyexpressed proteins (t•test;p-values<0.05, log2 foldchange <·1 or >1) between the two monoclonal cell lines C59 and Cl13. A-0: reactome pathways enriched for the comparison of C59 vs. C113 mock (uninfected) and C59 vs.C113Infected. Proteins separated Intoup-anddownregulated in eachcellline•C59Moup and C113 Mo up= upregulated in C59Mock or C113 Mock;C591nf up and C113 Inf up= upregulated in C59 Infected or C113 infected. Protein networ1< analysis was performed using the STRING platform and *H. sop/ens* as the reference proteome. Each protein Is assigned to the corresponding gene. Colors refer to functional clusters, which are connected by straight lines, genes of different clusters with a functional similarity are connected by dashed lines.Genes thatbelong to two functionalclusters are **marked with two different colors. Thename of the corresponding pathway of functional clusters islabeled inthe same color** as the genes. The PPIenrichment p-value is a statistical measure of the probability that a connection occurs randomly. **E,**G: heat maps for C59 and C113 were gnerated with thethree replicatesbased on significant differences (t-test, p<0.05) for the **z-scores usingOriginlab software. Protein network analysisof the clustersforproteinsdownregulated inC59infected(F) and** in C113 infected(H).

In addition, an upregulation of fatty acid beta-oxidation and a degradation of branched chain amino acids (BCAAs) such as valine, leucine and isoleucine was observed for the C113 clone (mock, Fig 6B). Both changes in the metabolism are described for mice, which are highly suspectable to IAV infection [56]. With regard to cell proliferation, an increase in beta oxidation can excert a positive or negative influence on cancer cells, as described previously [57]. Conversely, BCAA degradation has been shown to be associated with reduced growth through a reduced glucose uptake [58].

The data of the infected cells demonstrate that certain proteins related to infectious disease, stress response and TCA were upregulated in the C59 compared to the C113 clone (Fig 6C). A notable finding was the observation that the majority of proteins associated with stress response and infectious disease exhibited a significant overlap, suggesting a substantial interference between these two pathways. Given the broad nature of these two terms, a search was conducted for 15 genes related to infectious disease and stress response using the Reactome pathway database. This investigation revealed that three genes (NUP160, RPL30, DNAJC3) are directly linked to IAV RNA transcription and replication. Three other genes (PSMA5, PSMC4, RPN1) were linked to Wingless-related integration site (WNT) signaling, proliferation and DNA replication and another three genes (TUBB4B, CHMP4A, DYNC1H1) to vesicle-mediated transport. However, the association of the gene products with multiple signaling pathways complicates the evaluation of their impact (PSMA5 [59], PSMC4 [60], RPN1 [61]). For example, in one study it was shown, that activation of WNT signaling was positively correlated with IAV replication [62]. Liu et al. discussed that WNT signaling was downregulated during IAV infection [60]. Nevertheless, the observed hits for proliferation and DNA replication could be indicative of the tendency of the C59 clone for ongoing growth even during IAV infection.

In contrast, proteins of the C113 clone exhibited an increase in the expression of proteins related to membrane trafficking and the cell-mediated immunity. Four of the proteins associated with the latter were found to be directly linked to IAV NS1 (Fig 6D). In addition, previous studies demonstrated that the three nuclear pore complex proteins (NUP43, 58, 98) are involved in the transport between nucleus and cytoplasm by different viruses [63]-[65]. Apart from that, also antiviral pathways were triggered (Fig 6D). Specifically, RNAse L (RNASEL) and the RNA-activated kinase (EIF2AK2) have been identified as key players in antiviral responses, known to activate interferon pathways [66]-[68]. However, the impact of interferon signaling on IAV replication in MDCK cells is very low [69].

Membrane trafficking plays a critical role in the late stages of IAV infection, particularly during virus assembly and budding. During this step, the Ras-related protein Rab-11A (Rab11A), which has the highest number of cross-connections to other proteins in this study (Fig 5D), seems to be a key factor for the synthesis of new virus particles [70]-[71].

To facilitate a comparative analysis of the infected and the uninfected state (mock), heat maps were generated based on significant differences (t-test, p<0.05) in average z-values across the three replicates (Fig 6E, G). For the C59 clone, three clusters were identified, exhibiting a decrease in protein expression following infection. Based on all proteins from these clusters, a network can be derived that is mainly related to RNA metabolism (Fig 5F). It was evident that these proteins function as integral components of diverse processes, including ALK signaling, splicing, and virus replication (Supplementary Figure 2). For the C113 clone, apart from RNA processing, MET, mitogen-activated protein kinase signaling (MAPK) and TNF were also downregulated after infection (Fig 6G), all involved in antiviral effects and inhibition of IAV replication [72]-[74].

Next, we analyzed proteins that were exclusively expressed in infected cell clones or in mock samples (Fig 7A, B). For the C59 clone, only 26 proteins were present in all three infected replicates, whereas for C113, a total of 88 “unique” proteins were identified. These proteins were enriched and aligned to the corresponding pathways using the web-based portal Metascape [28]. For the C59 clone, the proteins were linked to carbohydrate metabolism, protein ubiquitination and the transport of amino acids and ions. In contrast, for C113, they were linked to different signaling processes such as apoptosis, TNF, VEGF and TOR signaling, as well as cytoskeleton and viral infection pathways. Notably, the lipid and carbohydrate metabolism were also affected by IAV repplication. Regarding the latter, an increase in the activity of the central carbohydrate and lipid metabolism is described to be benefical for efficient IAV replication since key proteins and structural parts of the virus are produced in higher quantities [75]. Especially during the late phase of IAV infection, TOR signaling and initiation of apoptosis by IAV are described to be beneficial for efficient virus assembly and release [76]-[77]. These discrepancies with the C59 clone may provide evidence to support the hypotheses that a key distinction between the two monoclonal cell lines lies is the efficient assembly and release of IAV particles.

**Figure 7:**
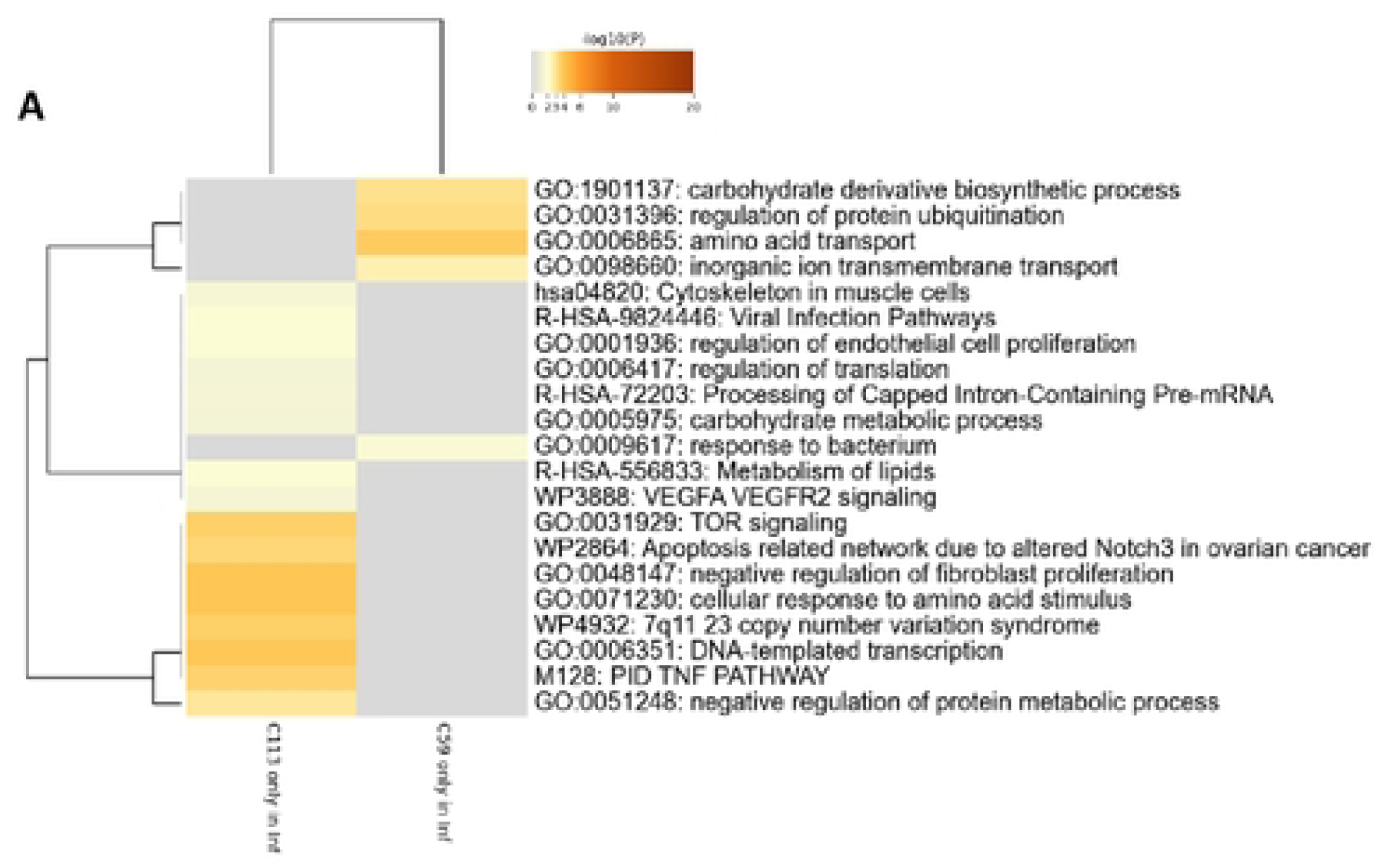
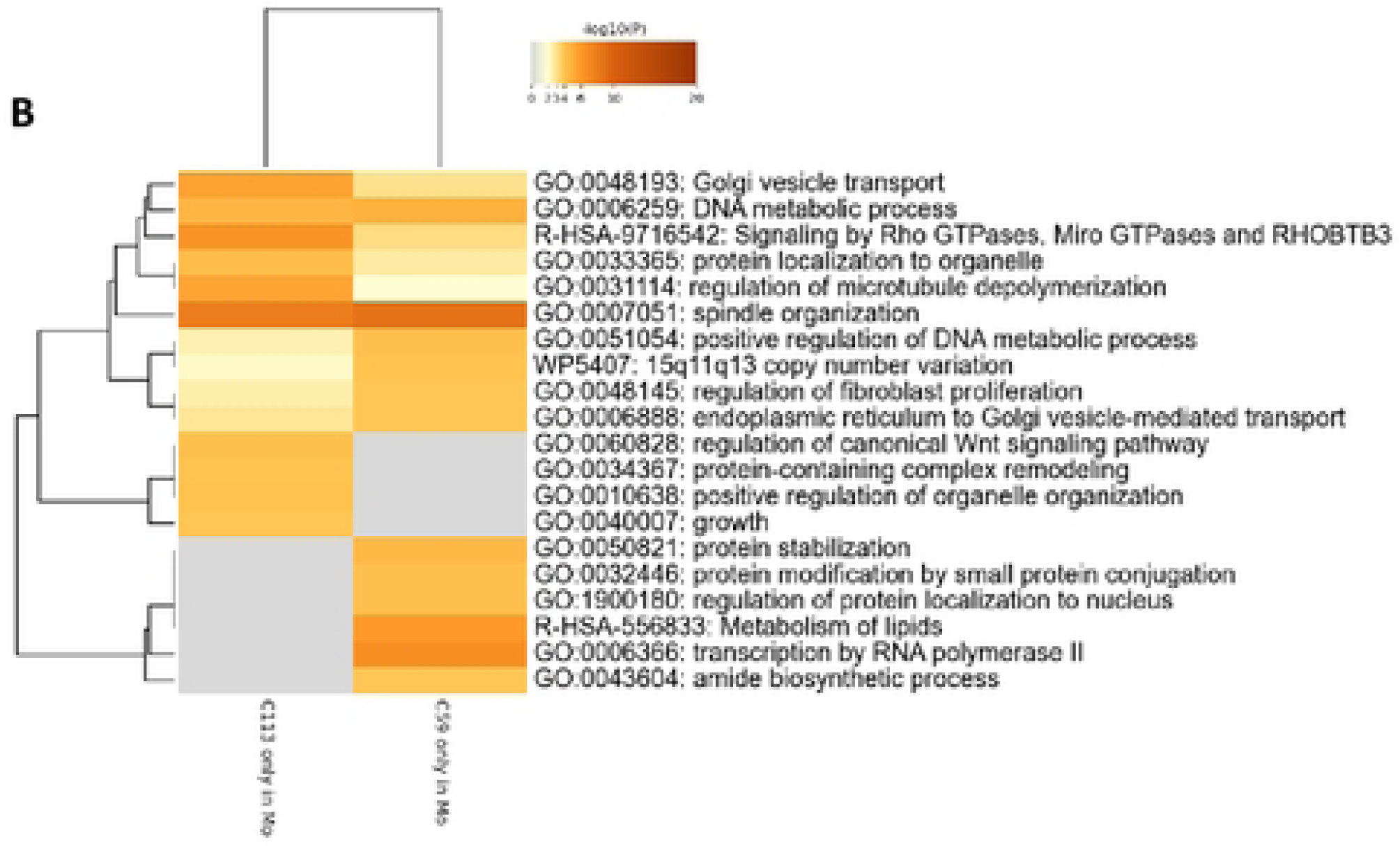
Metascape analysis of proteins detected in either A) infected CS9and Cll3 (CS9 only in Inf, C113 only in Inf), or **B)** only mock C59 and Cll3 (C59 only in Mo, C113 onlyin Mo) samples (n=3). Corresponding genes weremapped on the human genome and enriched asdescribed in Zhou et al., 2019 (28) using the default settings of the web tool.

A large number of proteins (379) were not identified in the infected samples of C59 compared to C113 (198 for C113 compared to C59). Many of these proteins are associated with related pathways, including Golgi transport, DNA metabolism and microtuble depolymerisation (Fig 7B). For C59, lipid metabolism and RNA polymerase II transcription appear to be inactivated following IAV infection, while for C113 proteins related to cell growth and regulation of WNT signaling were only present in the mock samples. In addition to the pivotal role of lipids in the replication of IAV, it is known that the cellular RNA polymerase II interacts with the RNA polymerase of IAV forming a complex for efficient replication of the virus [78]. The observed downregulation of this process may consequently impact the replication of the IAV genome, potentially resulting in reduced titers for the C59 clone. For C113, the shutdown of WNT signaling could limit IAV titer, as it is described that this signaling pathway can have pro-viral effects, especially in early stages of infection [63].

A comprehensive summary of all changes after infection of the two monoclonal cell lines is given in Figure 8. For the C113 clone, a number of antiviral intracellular signaling pathways were found to be downregulated during infection. In contrast, for C59, the JAK/STAT and the RAS pathways were upregulated. This could result in a limitation of the production and release of IAV particles in this cell line compared to C113. However, given that both cell lines produced significant amounts of IAV particles, it is evident that several virus-favouring factors were also activated. For the C59 clone, there is a high probability of increased viral compound production due to the highly active TCA and cholesterol synthesis pathways. Conversely, the C113 clone exhibited an increased presence of cellular factors that facilitate the efficient export of vRNPs and vmRNAs. It is conceivable that the obsereved differences in IAV titers may be attributable to an insufficent virus budding and release in the C59 clone in comparison to C113.

**Figure 8:**
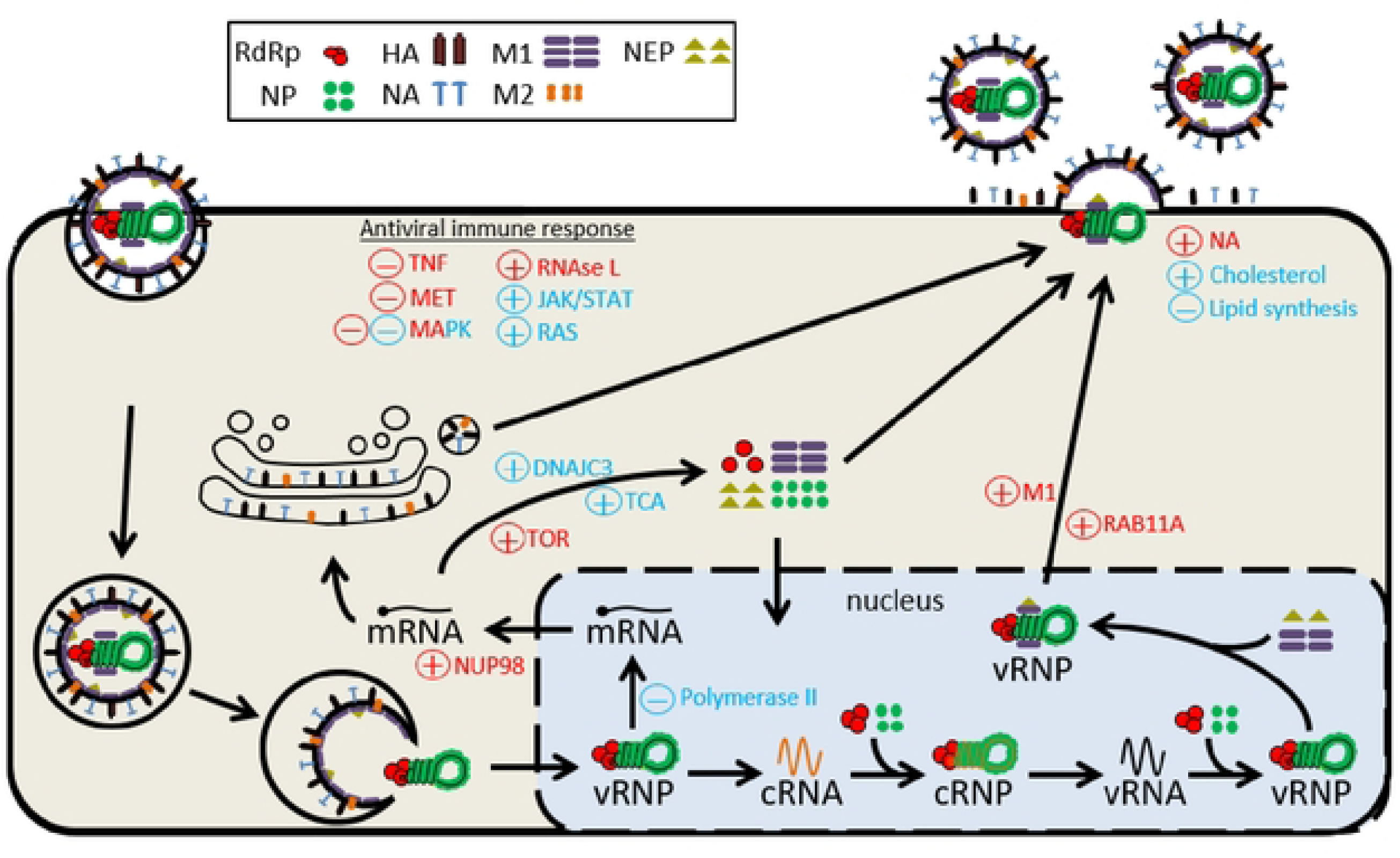
Summary of changes in the CS9 and C113 cell clones following IAV infection.+: upregulated; -: downregulated; red: changes in C113; blue: changes in C59. Adapted from (15]

Overall, a comprehensive analysis was conducted to identify key characteristics between a fast-growing and a high-producing clonal MDCK cell line. The results obtained facilitated the identification of potential bottlenecks of IAV replication and engendered a more profound comprehension of efficective IAV production in animal cell cultures. Increased membrane trafficking, activation of TOR signaling and apoptosis, and a reduction in antiviral signaling appear to be pivotal in fostering the high-yield production of IAV in MDCK cells.

## Author contributions

JK conducted the experiments and performed the analytics for the absolute quantification of the IAV and cell proteins. YG initiated the conceptualization of the study. JK, TZ, YG and DB planned the experiments. MW and TZ had key ideas in the analysis and visualization of the results and discussion and improved the written manuscript. MW wrote the python script for PCoA. PH adapted the DIA-PASEF method and performed the initial HCP analysis. UR provided funding, resources, and supported the interpretation and depiction of the results. JK wrote the paper. All authors read, commented, revised and approved the final version of the manuscript.

## Acknowledgements

The authors would like to thank Ilona Behrendt, Corina Siewert and Nancy Wynserski for their excellent technical support in cell cultivation and TCID_50_ assay execution. The authors would also like to thank the Sartorius Stedim Biotech GmbH for providing the clonal suspension cell lines and media.

## Ethical statement

This article does not contain any studies with human participants or animals performed by any of the authors.

## Data availability statement

Raw data files for MS measurements as well as processed spectra are uploaded to PanoramaWeb under the following link: https://panoramaweb.org/ozq7i2.url.

---

**Figure S1:**
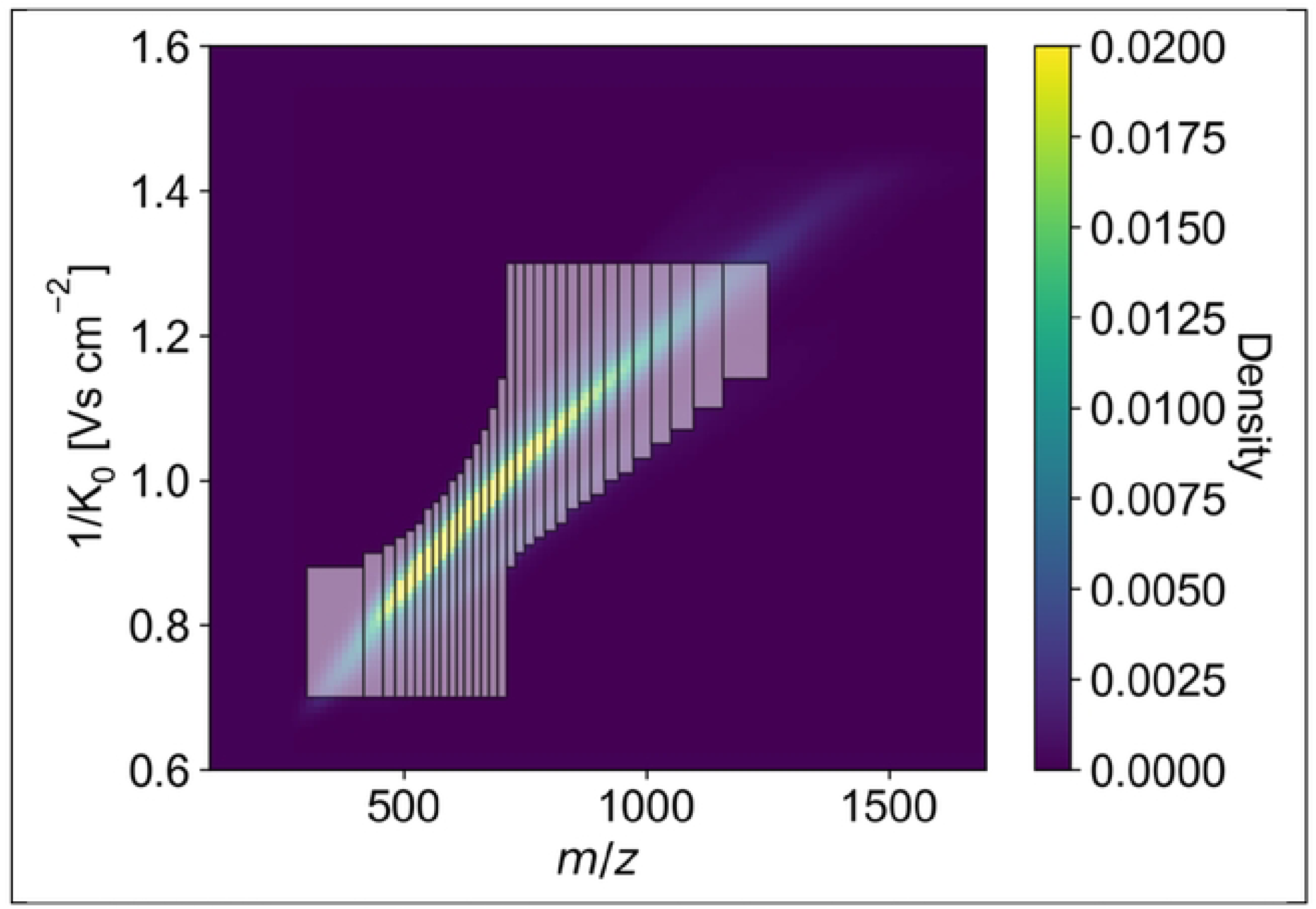
DIA-PASEF scan windows. Windows were selected based on ODA measurements using py diAID (Version 0.030) as describedin Skowronek et al. (25) resultingin a precursor coverage of 99.80%. Mobility (1/Ko) and m/z areas were adjusted in a manner that enabled the capture of the majority the ions, particularly those with ahigh density.

**Figure S2:**
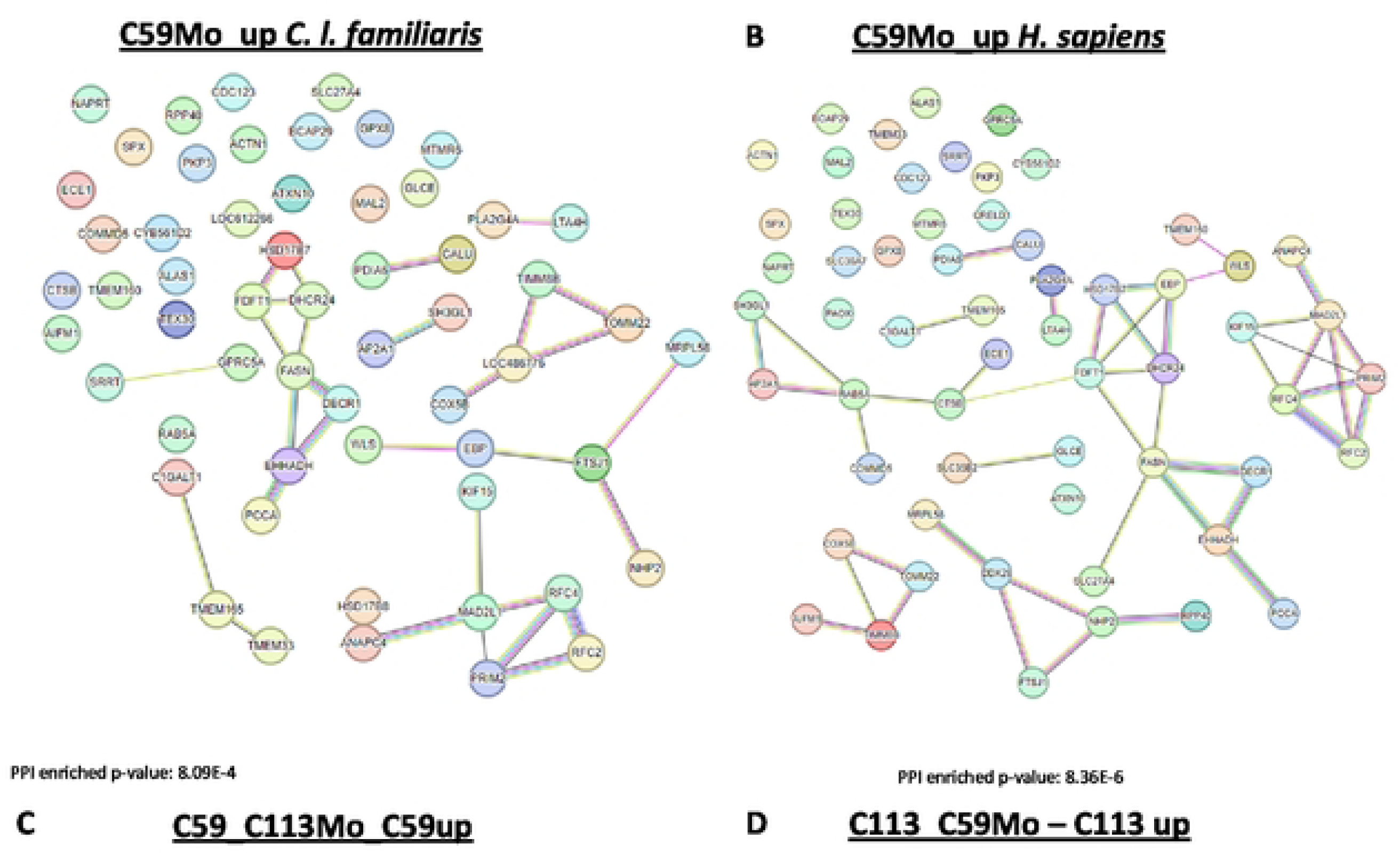
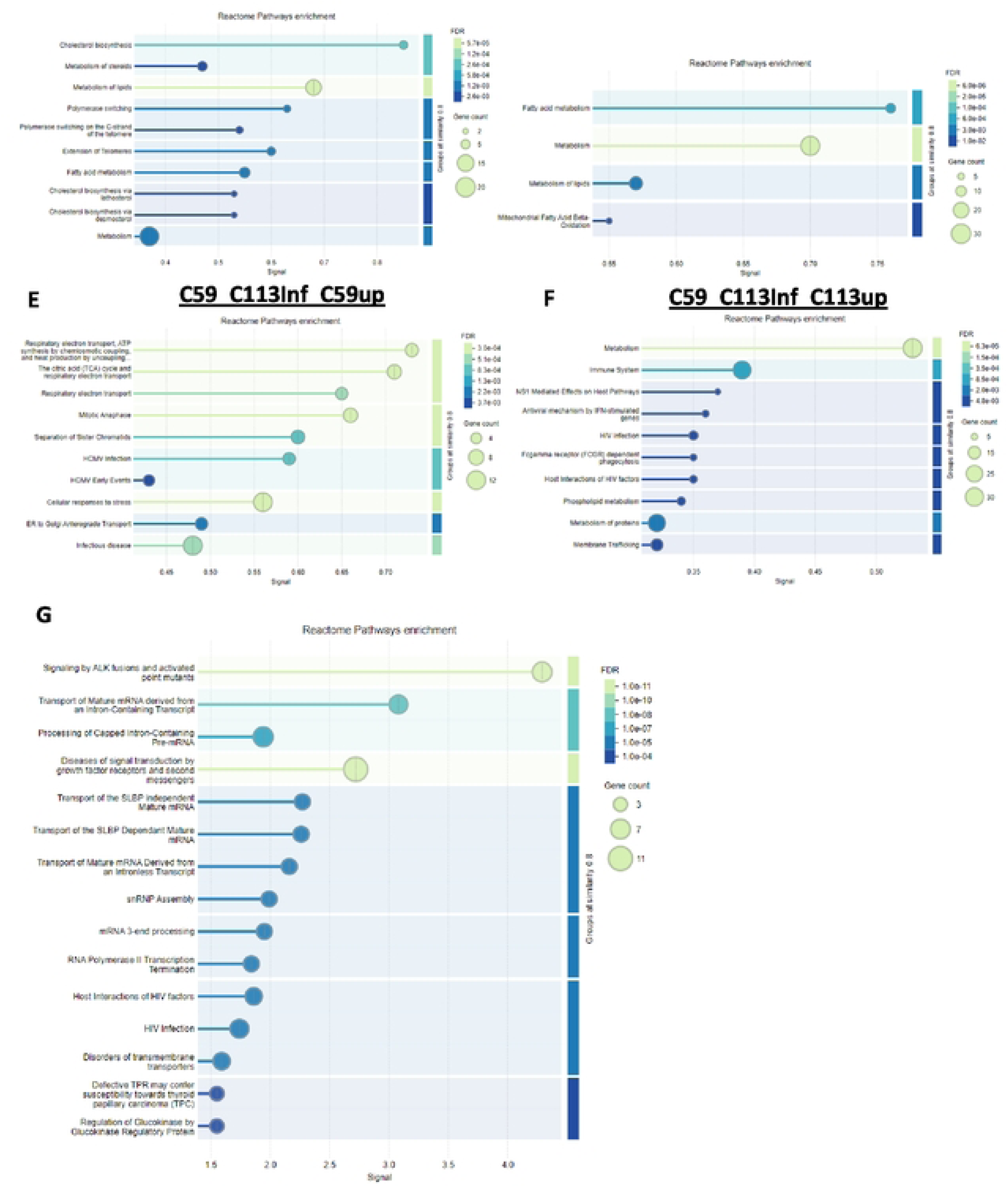
Protein network analysis of CS9infected and mock. Protein networks were generated using theplatform STRING for the comparison of CS9 and C113 Mock (C59 Mo,Cl13Mo) and infected (C59 Inf, C113 Inf). A, **8:** comparison of the two databases *C.*I.*familiaris* and*H. sapiens.* Each protein is assigned to the corresponding gene.ThePPIenrichment p•valueis a statistical measure of the probability thata connection occurs randomly. C,D,E,F:enriched reactorne pathways forC59 and Cl13 mock-infected and infected. G: enriched reactome pathways for the three dusters of downregulated proteins during infection in C59(referred to fig SE).

**Table S1:**
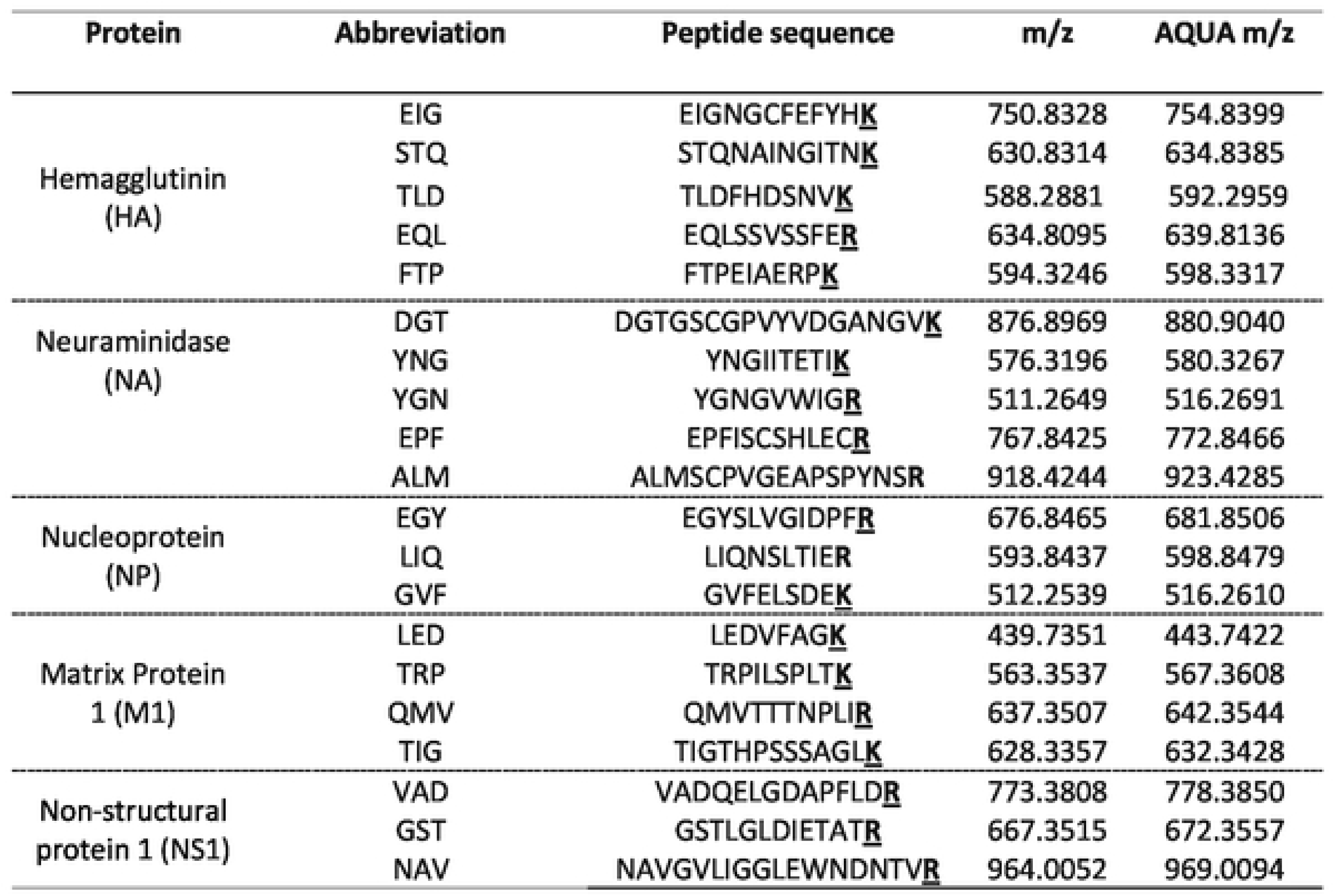
Target peptidesfor HA,NA. NP, Ml and NSl protein quantification. Underlined amino adds were C^13^ labelled and N^1^slabe.lied (adapted from(221).

**Table S2:** Errors of the absolute quantification of HA. NP, NA. Ml and NSl proteins. Technical error: percentage standard deviation of the abundance of all peptides divided through the absolute protein copy number (n=3); biological error: percentage standard deviation of three replicates.

|  | Technical error | Biological error |
| --- | --- | --- |
| Hemagglutinin (HA) | 30 % | 16 % |
| Nucleoprotein (NP) | 38 % | 21 % |
| Neuraminidase (NA) | 37 % | 26 % |
| Matrix protein 1 (M1) | 34 % | 22 % |
| Non-structural protein 1 (NS1) | 20 % | 14 % |
| Average | 32 % | 20 % |

**Table S3:** Limit of detection (LoD) andlimit of quantification (LoQ) for the HA and NP protein.

|  | LoD [copies/cell] | LoQ [copies/cell] |
| --- | --- | --- |
| Hemagglutinin (HA) | 5.8E+03 | 2.3E+04 |
| Nucleoprotein (NP) | 1.4E+04 | 5.6E+04 |

## References

1 Nicolson C, Major D, Wood JM, Robertson JS. Generation of influenza vaccine viruses on Vero cells by reverse genetics: an H5N1 candidate vaccine strain produced under a quality system. Vaccine. 2005; 23:2943–52. doi: 10.1016/j.vaccine.2004.08.054 PMID: 15780743.

2 Donis RO, Davis CT, Foust A, Hossain MJ, Johnson A, Klimov A, et al. Performance characteristics of qualified cell lines for isolation and propagation of influenza viruses for vaccine manufacturing. Vaccine. 2014; 32:6583–90. Epub 2014/06/24. doi: 10.1016/j.vaccine.2014.06.045 PMID: 24975811.

3 Bissinger T, Fritsch J, Mihut A, Wu Y, Liu X, Genzel Y, et al. Semi-perfusion cultures of suspension MDCK cells enable high cell concentrations and efficient influenza A virus production. Vaccine. 2019; 37:7003–10. Epub 2019/04/29. doi: 10.1016/j.vaccine.2019.04.054 PMID: 31047676.

4 Zinnecker T, Reichl U, Genzel Y (2024) Innovations in cell culture-based influenza vaccine manufacturing - from static cultures to high cell density cultivations. Hum Vaccin Immunother 20:2373521. doi: 10.1080/21645515.2024.2373521

5 Silva CAT, Kamen AA, Henry O. Recent advances and current challenges in process intensification of cell culture-based influenza virus vaccine manufacturing. Can J Chem Eng. 2021; 99:2525–35. doi: 10.1002/cjce.24197.

6 Zinnecker T, Badri N, Araujo D, Thiele K, Reichl U, Genzel Y. From single-cell cloning to high-yield influenza virus production - implementing advanced technologies in vaccine process development. Eng Life Sci. 2024; 24:2300245. Epub 2024/02/18. doi: 10.1002/elsc.202300245 PMID: 38584687.

7 Vlecken DHW, Pelgrim RPM, Ruminski S, Bakker WAM, van der Pol LA. Comparison of initial feasibility of host cell lines for viral vaccine production. J Virol Methods. 2013; 193:28–41. Epub 2013/05/14. doi: 10.1016/j.jviromet.2013.04.020 PMID: 23684847.

8 Genzel Y. Designing cell lines for viral vaccine production: Where do we stand. Biotechnol J. 2015; 10:728–40. Epub 2015/04/22. doi: 10.1002/biot.201400388 PMID: 25903999.

9 Meguro H, Bryant JD, Torrence AE, Wright PF. Canine kidney cell line for isolation of respiratory viruses. J Clin Microbiol. 1979; 9:175–9. doi: 10.1128/jcm.9.2.175-179.1979 PMID: 219021.

10 Liu Y, Childs RA, Matrosovich T, Wharton S, Palma AS, Chai W, et al. Altered receptor specificity and cell tropism of D222G hemagglutinin mutants isolated from fatal cases of pandemic A(H1N1) 2009 influenza virus. J Virol. 2010; 84:12069–74. Epub 2010/09/08. doi: 10.1128/jvi.01639-10 PMID: 20826688.

11 Peschel B, Frentzel S, Laske T, Genzel Y, Reichl U. Comparison of influenza virus yields and apoptosis-induction in an adherent and a suspension MDCK cell line. Vaccine. 2013; 31:5693–9. Epub 2013/10/08. doi: 10.1016/j.vaccine.2013.09.051 PMID: 24113260.

12 Tapia F, Vázquez-Ramírez D, Genzel Y, Reichl U. Bioreactors for high cell density and continuous multi-stage cultivations: options for process intensification in cell culture-based viral vaccine production. Appl Microbiol Biotechnol. 2016; 100:2121–32. Epub 2016/01/13. doi: 10.1007/s00253-015-7267-9 PMID: 26758296.

13 Liu J, Shi X, Schwartz R, Kemble G. Use of MDCK cells for production of live attenuated influenza vaccine. Vaccine. 2009; 27:6460–3. Epub 2009/06/24. doi: 10.1016/j.vaccine.2009.06.024 PMID: 19559113.

14 Lugovtsev VY, Melnyk D, Weir JP. Heterogeneity of the MDCK cell line and its applicability for influenza virus research. PLoS One. 2013; 8:e75014. Epub 2013/09/13. doi: 10.1371/journal.pone.0075014 PMID: 24058646.

15 Heldt FS, Kupke SY, Dorl S, Reichl U, Frensing T. Single-cell analysis and stochastic modelling unveil large cell-to-cell variability in influenza A virus infection. Nat Commun. 2015; 6:8938. Epub 2015/11/20. doi: 10.1038/ncomms9938 PMID: 26586423.

16 Kluge S, Benndorf D, Genzel Y, Scharfenberg K, Rapp E, Reichl U. Monitoring changes in proteome during stepwise adaptation of a MDCK cell line from adherence to growth in suspension. Vaccine. 2015; 33:4269–80. Epub 2015/04/16. doi: 10.1016/j.vaccine.2015.02.077 PMID: 25891398.

17 Pralow A, Hoffmann M, Nguyen-Khuong T, Pioch M, Hennig R, Genzel Y, et al. Comprehensive N-glycosylation analysis of the influenza A virus proteins HA and NA from adherent and suspension MDCK cells. FEBS J. 2021; 288:4869–91. Epub 2021/03/15. doi: 10.1111/febs.15787 PMID: 33629527.

18 Pan S, Aebersold R, Chen R, Rush J, Goodlett DR, McIntosh MW, et al. Mass spectrometry based targeted protein quantification: methods and applications. J Proteome Res. 2009; 8:787–97. doi: 10.1021/pr800538n PMID: 19105742.

19 Sahoo A, Tsukiadate T, Lin B-R, Kotzbauer E, Houser J, Patel M, et al. Proteomics Reveals Distinctive Host Cell Protein Expression Patterns in Fed-Batch and Perfusion Cell Culture Processes. Biotechnol J. 2025; 20:e202400567. doi: 10.1002/biot.202400567 PMID: 39834099.

20 Kalbfuss B, Knöchlein A, Kröber T, Reichl U. Monitoring influenza virus content in vaccine production: precise assays for the quantitation of hemagglutination and neuraminidase activity. Biologicals. 2008; 36:145–61. doi: 10.1016/j.biologicals.2007.10.002 PMID: 18561375.

21. Genzel Y, Reichl U. Vaccine Production. In: Walker JM, Pörtner R, editors. Animal Cell Biotechnology. Totowa, NJ: Humana Press; 2007. pp. 457–73.

22 Küchler J, Opitz P, Jordan I, Genzel Y, Benndorf D, Reichl U. Quantification of intracellular influenza A virus protein dynamics in different host cells after seed virus adaptation. Appl Microbiol Biotechnol. 2025; 109:74. Epub 2025/03/24. doi: 10.1007/s00253-025-13423-3 PMID: 40126655.

23 Heyer R, Schallert K, Büdel A, Zoun R, Dorl S, Behne A, et al. A Robust and Universal Metaproteomics Workflow for Research Studies and Routine Diagnostics Within 24 h Using Phenol Extraction, FASP Digest, and the MetaProteomeAnalyzer. Front Microbiol. 2019; 10:1883. Epub 2019/08/16. doi: 10.3389/fmicb.2019.01883 PMID: 31474963.

24 Rüdiger D, Piasecka J, Küchler J, Pontes C, Laske T, Kupke SY, et al. Mathematical model calibrated to in vitro data predicts mechanisms of antiviral action of the influenza defective interfering particle "OP7". iScience. 2024; 27:109421. Epub 2024/03/05. doi: 10.1016/j.isci.2024.109421 PMID: 38523782.

25 Skowronek P, Wallmann G, Wahle M, Willems S, Mann M. An accessible workflow for high-sensitivity proteomics using parallel accumulation-serial fragmentation (PASEF). Nat Protoc. 2025. Epub 2025/01/17. doi: 10.1038/s41596-024-01104-w PMID: 39825144.

26 Demichev V, Szyrwiel L, Yu F, Teo GC, Rosenberger G, Niewienda A, et al. dia-PASEF data analysis using FragPipe and DIA-NN for deep proteomics of low sample amounts. Nat Commun. 2022; 13:3944. Epub 2022/07/08. doi: 10.1038/s41467-022-31492-0 PMID: 35803928.

27 Schiebenhoefer H, Schallert K, Renard BY, Trappe K, Schmid E, Benndorf D, Riedel K, Muth T, Fuchs S (2020) A complete and flexible workflow for metaproteomics data analysis based on MetaProteomeAnalyzer and Prophane. Nat Protoc 15:3212–3239. doi: 10.1038/s41596-020-0368-7

28 Szklarczyk D, Kirsch R, Koutrouli M, Nastou K, Mehryary F, Hachilif R, et al. The STRING database in 2023: protein-protein association networks and functional enrichment analyses for any sequenced genome of interest. Nucleic Acids Res. 2023; 51:D638–D646. doi: 10.1093/nar/gkac1000 PMID: 36370105.

29. Zhou Y, Zhou B, Pache L, Chang M, Khodabakhshi AH, Tanaseichuk O, et al. Metascape provides a biologist-oriented resource for the analysis of systems-level datasets. Nat Commun. 2019; 10:1523. Epub 2019/04/03. doi: 10.1038/s41467-019-09234-6 PMID: 30944313.

30. Le Grimellec C, Lesniewska E, Cachia C, Schreiber JP, Fornel F de, Goudonnet JP (1994) Imaging of the membrane surface of MDCK cells by atomic force microscopy. Biophys J 67:36–41. doi: 10.1016/S0006-3495(94)80490-4

31 Lange K (2002) Role of microvillar cell surfaces in the regulation of glucose uptake and organization of energy metabolism. Am J Physiol Cell Physiol 282:C1–26. doi: 10.1152/ajpcell.2002.282.1.C1

32 Kolesnikova L, Heck S, Matrosovich T, Klenk H-D, Becker S, Matrosovich M (2013) Influenza virus budding from the tips of cellular microvilli in differentiated human airway epithelial cells. J Gen Virol 94:971–976. doi: 10.1099/vir.0.049239-0

33 Jagadesh A, Salam AAA, Mudgal PP, Arunkumar G. Influenza virus neuraminidase (NA): a target for antivirals and vaccines. Arch Virol. 2016; 161:2087–94. Epub 2016/06/02. doi: 10.1007/s00705-016-2907-7 PMID: 27255748.

34 Wagner R, Matrosovich M, Klenk H-D (2002) Functional balance between haemagglutinin and neuraminidase in influenza virus infections. Rev Med Virol 12:159–166. doi: 10.1002/rmv.352

35 Ali A, Avalos RT, Ponimaskin E, Nayak DP (2000) Influenza virus assembly: effect of influenza virus glycoproteins on the membrane association of M1 protein. J Virol 74:8709–8719. doi: 10.1128/jvi.74.18.8709-8719.2000

36 Qi F, Li Y, Yang X, Wu Y-P, Lin L-J, Liu X-M. Significance of alternative splicing in cancer cells. Chin Med J (Engl). 2020; 133:221–8. doi: 10.1097/CM9.0000000000000542 PMID: 31764175.

37 van den Broeke C, Jacob T, Favoreel HW. Rho’ing in and out of cells: viral interactions with Rho GTPase signaling. Small GTPases. 2014; 5:e28318. Epub 2014/03/24. doi: 10.4161/sgtp.28318 PMID: 24691164.

38 Crosas-Molist E, Samain R, Kohlhammer L, Orgaz JL, George SL, Maiques O, et al. Rho GTPase signaling in cancer progression and dissemination. Physiol Rev. 2022; 102:455–510. Epub 2021/09/20. doi: 10.1152/physrev.00045.2020 PMID: 34541899.

39 Michalik L, Wahli W. PPARs Mediate Lipid Signaling in Inflammation and Cancer. PPAR Res. 2008; 2008:134059. Epub 2008/12/21. doi: 10.1155/2008/134059 PMID: 19125181.

40 Fukao T, Mitchell GA, Song XQ, Nakamura H, Kassovska-Bratinova S, Orii KE, et al. Succinyl-CoA:3- ketoacid CoA transferase (SCOT): cloning of the human SCOT gene, tertiary structural modeling of the human SCOT monomer, and characterization of three pathogenic mutations. Genomics. 2000; 68:144– 51. doi: 10.1006/geno.2000.6282 PMID: 10964512.

41 Zhang S, Xie C. The role of OXCT1 in the pathogenesis of cancer as a rate-limiting enzyme of ketone body metabolism. Life Sci. 2017; 183:110–5. Epub 2017/07/04. doi: 10.1016/j.lfs.2017.07.003 PMID: 28684065.

42 Mass E, Wachten D, Aschenbrenner AC, Voelzmann A, Hoch M. Murine Creld1 controls cardiac development through activation of calcineurin/NFATc1 signaling. Dev Cell. 2014; 28:711–26. doi: 10.1016/j.devcel.2014.02.012 PMID: 24697899.

43 Fang M, Tang X, Zhang J, Liao Z, Wang G, Cheng R, et al. An inhibitor of leukotriene-A4 hydrolase from bat salivary glands facilitates virus infection. Proc Natl Acad Sci U S A. 2022; 119:e2110647119. Epub 2022/03/01. doi: 10.1073/pnas.2110647119 PMID: 35238649.

44 Quistgaard EM. BAP31: Physiological functions and roles in disease. Biochimie. 2021; 186:105–29. Epub 2021/04/28. doi: 10.1016/j.biochi.2021.04.008 PMID: 33930507.

45 Daugherty M, Polanuyer B, Farrell M, Scholle M, Lykidis A, Crécy-Lagard V de, et al. Complete reconstitution of the human coenzyme A biosynthetic pathway via comparative genomics. J Biol Chem. 2002; 277:21431–9. Epub 2002/03/28. doi: 10.1074/jbc.M201708200 PMID: 11923312.

46 Breus OS, Nemazanyy IO, Gout IT, Filonenko VV, Panasyuk GG. CoA Synthase influences adherence-independent growth and survival of mammalian cells in vitro. Biopolym Cell. 2009; 25:384–9. doi: 10.7124/bc.0007F0.

47 Ehrhardt C, Ludwig S. A new player in a deadly game: influenza viruses and the PI3K/Akt signalling pathway. Cell Microbiol. 2009; 11:863–71. Epub 2009/03/12. doi: 10.1111/j.1462-5822.2009.01309.x PMID: 19290913.

48 Hu X, Han Y, Liu J, Wang H, Tian Z, Zhang X, et al. CTP synthase 2 predicts inferior survival and mediates DNA damage response via interacting with BRCA1 in chronic lymphocytic leukemia. Exp Hematol Oncol. 2023; 12:6. Epub 2023/01/12. doi: 10.1186/s40164-022-00364-0 PMID: 36635772.

49 Lee K-W, Okot-Kotber C, LaComb JF, Bogenhagen DF. Mitochondrial ribosomal RNA (rRNA) methyltransferase family members are positioned to modify nascent rRNA in foci near the mitochondrial DNA nucleoid. J Biol Chem. 2013; 288:31386–99. Epub 2013/09/13. doi: 10.1074/jbc.M113.515692 PMID: 24036117.

50 Sieczkarski SB, Whittaker GR. Dissecting virus entry via endocytosis. J Gen Virol. 2002; 83:1535–45. doi: 10.1099/0022-1317-83-7-1535 PMID: 12075072.

51 Sun X, Whittaker GR. Role for influenza virus envelope cholesterol in virus entry and infection. J Virol. 2003; 77:12543–51. doi: 10.1128/jvi.77.23.12543-12551.2003 PMID: 14610177.

52 Tüzmen Ş, Hostetter G, Watanabe A, Ekmekçi C, Carrigan PE, Shechter I, et al. Characterization of farnesyl diphosphate farnesyl transferase 1 (FDFT1) expression in cancer. Per Med. 2019; 16:51–65. Epub 2018/11/23. doi: 10.2217/pme-2016-0058 PMID: 30468409.

53 Assefi M, Bijan Rostami R, Ebrahimi M, Altafi M, Tehrany PM, Zaidan HK, et al. Potential use of the cholesterol transfer inhibitor U18666A as an antiviral drug for research on various viral infections. Microb Pathog. 2023; 179:106096. Epub 2023/04/01. doi: 10.1016/j.micpath.2023.106096 PMID: 37011734.

54 Zhao L, Shen G, Luo J, Zhang Y, Yao Y, Cui L, et al. Effects of Steroidal Compounds on Viruses. Viral Immunol. 2025; 38:44–52. Epub 2025/02/06. doi: 10.1089/vim.2024.0011 PMID: 39912855.

55 Liu J, Geng X, Hou J, Wu G. New insights into M1/M2 macrophages: key modulators in cancer progression. Cancer Cell Int. 2021; 21:389. Epub 2021/07/21. doi: 10.1186/s12935-021-02089-2 PMID: 34289846.

56 van Liempd S, Cabrera D, Pilzner C, Kollmus H, Schughart K, Falcón-Pérez JM. Impaired beta-oxidation increases vulnerability to influenza A infection. J Biol Chem. 2021; 297:101298. Epub 2021/10/09. doi: 10.1016/j.jbc.2021.101298 PMID: 34637789.

57 Currie E, Schulze A, Zechner R, Walther TC, Farese RV. Cellular fatty acid metabolism and cancer. Cell Metab. 2013; 18:153–61. Epub 2013/06/20. doi: 10.1016/j.cmet.2013.05.017 PMID: 23791484.

58 Shao D, Villet O, Zhang Z, Choi SW, Yan J, Ritterhoff J, et al. Glucose promotes cell growth by suppressing branched-chain amino acid degradation. Nat Commun. 2018; 9:2935. Epub 2018/07/26. doi: 10.1038/s41467-018-05362-7 PMID: 30050148.

59 Lu F, Zhou J, Chen Q, Zhu J, Zheng X, Fang N, et al. PSMA5 contributes to progression of lung adenocarcinoma in association with the JAK/STAT pathway. Carcinogenesis. 2022; 43:624–34. doi: 10.1093/carcin/bgac046 PMID: 35605971.

60 Liu K, Zhang S, Gong Y, Zhu P, Shen W, Zhang Q. PSMC4 promotes prostate carcinoma progression by regulating the CBX3-EGFR-PI3K-AKT-mTOR pathway. J Cell Mol Med. 2023; 27:2437–47. Epub 2023/07/12. doi: 10.1111/jcmm.17832 PMID: 37436074.

61 Luo P, Li Z, He H, Tang Y, Zeng L, Luo L, et al. Exploring the impact of RPN1 on tumorigenesis and immune response in cancer. FASEB J. 2025; 39:e70345. doi: 10.1096/fj.202401088R PMID: 40079196.

62 More S, Yang X, Zhu Z, Bamunuarachchi G, Guo Y, Huang C, et al. Regulation of influenza virus replication by Wnt/β-catenin signaling. PLoS One. 2018; 13:e0191010. Epub 2018/01/11. doi: 10.1371/journal.pone.0191010 PMID: 29324866.

63 Ling Y-H, Wang H, Han M-Q, Di Wang, Hu Y-X, Zhou K, et al. Nucleoporin 85 interacts with influenza A virus PB1 and PB2 to promote its replication by facilitating nuclear import of ribonucleoprotein. Front Microbiol. 2022; 13:895779. Epub 2022/08/16. doi: 10.3389/fmicb.2022.895779 PMID: 36051755.

64 Khanna M, Sharma K, Saxena SK, Sharma JG, Rajput R, Kumar B. Unravelling the interaction between Influenza virus and the nuclear pore complex: insights into viral replication and host immune response. Virusdisease. 2024; 35:231–42. Epub 2024/07/18. doi: 10.1007/s13337-024-00879-6 PMID: 39071870.

65 Zhang X, Lim K, Qiu Y, Hazawa M, Wong RW. Strategies for the Viral Exploitation of Nuclear Pore Transport Pathways. Viruses. 2025; 17. Epub 2025/01/23. doi: 10.3390/v17020151 PMID: 40006906.

66 Hatada E, Saito S, Fukuda R. Mutant influenza viruses with a defective NS1 protein cannot block the activation of PKR in infected cells. J Virol. 1999; 73:2425–33. doi: 10.1128/jvi.73.3.2425-2433.1999 PMID: 9971827.

67 Cooper DA, Banerjee S, Chakrabarti A, García-Sastre A, Hesselberth JR, Silverman RH, et al. RNase L targets distinct sites in influenza A virus RNAs. J Virol. 2015; 89:2764–76. Epub 2014/12/24. doi: 10.1128/jvi.02953-14 PMID: 25540362.

68 Le Y, Zhang J, Gong Z, Zhang Z, Nian X, Li X, et al. TRAF3 deficiency in MDCK cells improved sensitivity to the influenza A virus. Heliyon. 2023; 9:e19246. Epub 2023/08/21. doi: 10.1016/j.heliyon.2023.e19246 PMID: 37681145.

69 Seitz C, Isken B, Heynisch B, Rettkowski M, Frensing T, Reichl U. Trypsin promotes efficient influenza vaccine production in MDCK cells by interfering with the antiviral host response. Appl Microbiol Biotechnol. 2012; 93:601–11. Epub 2011/09/14. doi: 10.1007/s00253-011-3569-8 PMID: 21915610.

70 Eisfeld AJ, Kawakami E, Watanabe T, Neumann G, Kawaoka Y. RAB11A is essential for transport of the influenza virus genome to the plasma membrane. J Virol. 2011; 85:6117–26. Epub 2011/04/27. doi: 10.1128/jvi.00378-11 PMID: 21525351.

71 Dou D, Revol R, Östbye H, Wang H, Daniels R. Influenza A Virus Cell Entry, Replication, Virion Assembly and Movement. Front Immunol. 2018; 9:1581. Epub 2018/07/20. doi: 10.3389/fimmu.2018.01581 PMID: 30079062.

72 Matikainen S, Sirén J, Tissari J, Veckman V, Pirhonen J, Severa M, et al. Tumor necrosis factor alpha enhances influenza A virus-induced expression of antiviral cytokines by activating RIG-I gene expression. J Virol. 2006; 80:3515–22. doi: 10.1128/jvi.80.7.3515-3522.2006 PMID: 16537619.

73 Yu J, Sun X, Goie JYG, Zhang Y. Regulation of Host Immune Responses against Influenza A Virus Infection by Mitogen-Activated Protein Kinases (MAPKs). Microorganisms. 2020; 8. Epub 2020/07/17. doi: 10.3390/microorganisms8071067 PMID: 32709018.

74 Imamura R, Sato H, Chih-Cheng Voon D, Shirasaki T, Honda M, Kurachi M, et al. Met receptor is essential for MAVS-mediated antiviral innate immunity in epithelial cells independent of its kinase activity. Proc Natl Acad Sci U S A. 2023; 120:e2307318120. Epub 2023/09/25. doi: 10.1073/pnas.2307318120 PMID: 37748074.

75 Keshavarz M, Solaymani-Mohammadi F, Namdari H, Arjeini Y, Mousavi MJ, Rezaei F. Metabolic host response and therapeutic approaches to influenza infection. Cell Mol Biol Lett. 2020; 25:15. Epub 2020/03/05. doi: 10.1186/s11658-020-00211-2 PMID: 32161622.

76 Kuss-Duerkop SK, Wang J, Mena I, White K, Metreveli G, Sakthivel R, et al. Influenza virus differentially activates mTORC1 and mTORC2 signaling to maximize late stage replication. PLoS Pathog. 2017; 13:e1006635. Epub 2017/09/27. doi: 10.1371/journal.ppat.1006635 PMID: 28953980.

77 Ampomah PB, Lim LHK. Influenza A virus-induced apoptosis and virus propagation. Apoptosis. 2020; 25:1–11. doi: 10.1007/s10495-019-01575-3 PMID: 31667646.

78 Engelhardt OG, Smith M, Fodor E. Association of the influenza A virus RNA-dependent RNA polymerase with cellular RNA polymerase II. J Virol. 2005; 79:5812–8. doi: 10.1128/jvi.79.9.5812-5818.2005 PMID: 15827195.

